# DNA Framework Nanoreactor for Programmable Membrane Fusion

**DOI:** 10.1101/2025.10.28.685246

**Authors:** Qian Shi, Qiulan Yang, Fan Li, Min Bao, Shengwen Wang, Kui Huang, Liu Jiayi, Yunyun Wang, Yuanfang Chen, Yuhe Renee Yang, Xin Bian, Zhenyong Wu, Yang Yang

## Abstract

Membrane fusion is a fundamental yet transient process that has long resisted direct structural and kinetic dissection. Here we introduce a DNA framework vesicle (DFV) nanoreactor that transforms this elusive biological phenomenon into a programmable, visualizable, and quantifiable process-a nanoreaction confined in space yet extended in time. By embedding lipid membranes and SNARE proteins within precisely defined DNA apertures, DFVs convert stochastic vesicle collisions into geometry-controlled fusion events with tunable kinetics. Cryo-electron microscopy resolves a complete sequence of six intermediates, revealing how nanoscale confinement reshapes the energetic landscape of bilayer merger. Quantitative fluorescence and nano-flow cytometry further establish a direct link between spatial design and fusion probability. Extending this concept to living cells, DFVs enable controllable membrane fusion and augmentation on VAMP2-expressing membranes, achieving direct cytosolic delivery of functional siRNA via fusion-driven transfer rather than endocytosis. This framework bridges structural precision and functional mimicry, offering a unified platform to reconstruct, quantify, and harness membrane fusion as a programmable process for synthetic biology and nanomedicine.

## Introduction

Membrane fusion is a fundamental process in cell biology, essential for biomolecular transport, organelle maintenance, and various signaling events. It underlies critical biological functions such as neurotransmitter release, endocytosis, and the entry of enveloped viruses into host cells (*1–4*). This process proceeds through a series of tightly orchestrated steps—membrane apposition, protein-mediated docking, lipid bilayer mixing, and the formation and dilation of a fusion pore—that ultimately merge two separate lipid membranes into one continuous structure (*2*). Among the molecular drivers of membrane fusion, SNARE (Soluble N-ethylmaleimide-sensitive factor Attachment protein REceptor) proteins have been the most extensively studied. They form trans-complexes bridging opposing membranes and drive fusion by progressive “zippering,” which destabilizes local membrane structure (*5*). Yet, the short-lived and dynamic nature of fusion intermediates—particularly pore formation and dilation—has made direct visualization and mechanistic dissection exceedingly difficult (*6*).

Traditional experimental models, including native vesicles and synthetic proteoliposomes (*7, 8*), typically rely on ensemble fluorescence-based assays such as lipid mixing (FRET) or content dequenching to infer fusion activity (*6, 9*). However, these methods lack precise control over membrane geometry and protein positioning. The inherent variability in vesicle size, the stochastic nature of vesicle collisions, and the presence of multiple, unsynchronized fusion events further complicate the interpretation of intermediate states and energy landscapes (*2, 6*).

DNA nanotechnology provides a versatile toolbox to address the spatial and organizational challenges inherent in membrane fusion studies. In earlier work, Lin and coworkers developed a DNA origami nano-ring system that served as a structural exoskeleton to direct the formation of liposomes within a confined cavity via lipid mixing and detergent removal (*10*). This approach allowed the generation of liposomes with controlled size and shape, and enabled programmable vesicle positioning, protein stoichiometry adjustment, and membrane tethering (*11–13*). However, the exposed hemispherical surface of liposomes in this ring-based system left the actual fusion interface—particularly the nascent fusion pore—insufficiently constrained. As a result, the dynamic and localized nature of pore formation and expansion remained beyond the reach of structural capture. These limitations highlighted a critical need: to confine the membrane contact zone to a well-defined nanoscale aperture that could isolate and stabilize fusion intermediates.

To overcome this, we designed a DNA soccer-ball framework (DSF)—a truncated icosahedral origami structure—to template uniform liposomes and mediate defined vesicle–vesicle interactions. These DNA Framework Vesicles (DFVs) can be tethered via complementary strands at specific windows, confining the fusion interface within precisely aligned windows. This geometric confinement effectively trades physical space for temporal resolution, stabilizing transient intermediates thus providing a controlled setting to dissect SNAREs mediated fusion events.

We demonstrate that these small, window-defined contact zones can stabilize key intermediates such as hemifusion diaphragms and nascent pores, thereby enabling their direct structural capture via cryo-EM. In addition to precise parameter modulation — including membrane composition, window geometry, and inter-frame distance — this framework-based platform supports the modular assembly of heterotypic and higher-order structures (**Figure 1a**). We further show that DFVs can undergo SNARE-mediated membrane fusion directly on the surface of living cells, facilitating cytosolic delivery of siRNA via a fusion-driven mechanism. Together, these results establish DNA framework vesicles as a programmable and versatile system for high-resolution mechanistic studies and targeted cargo delivery, with broad potential in membrane engineering, synthetic cell design, and therapeutic applications.

**Figure 1.**
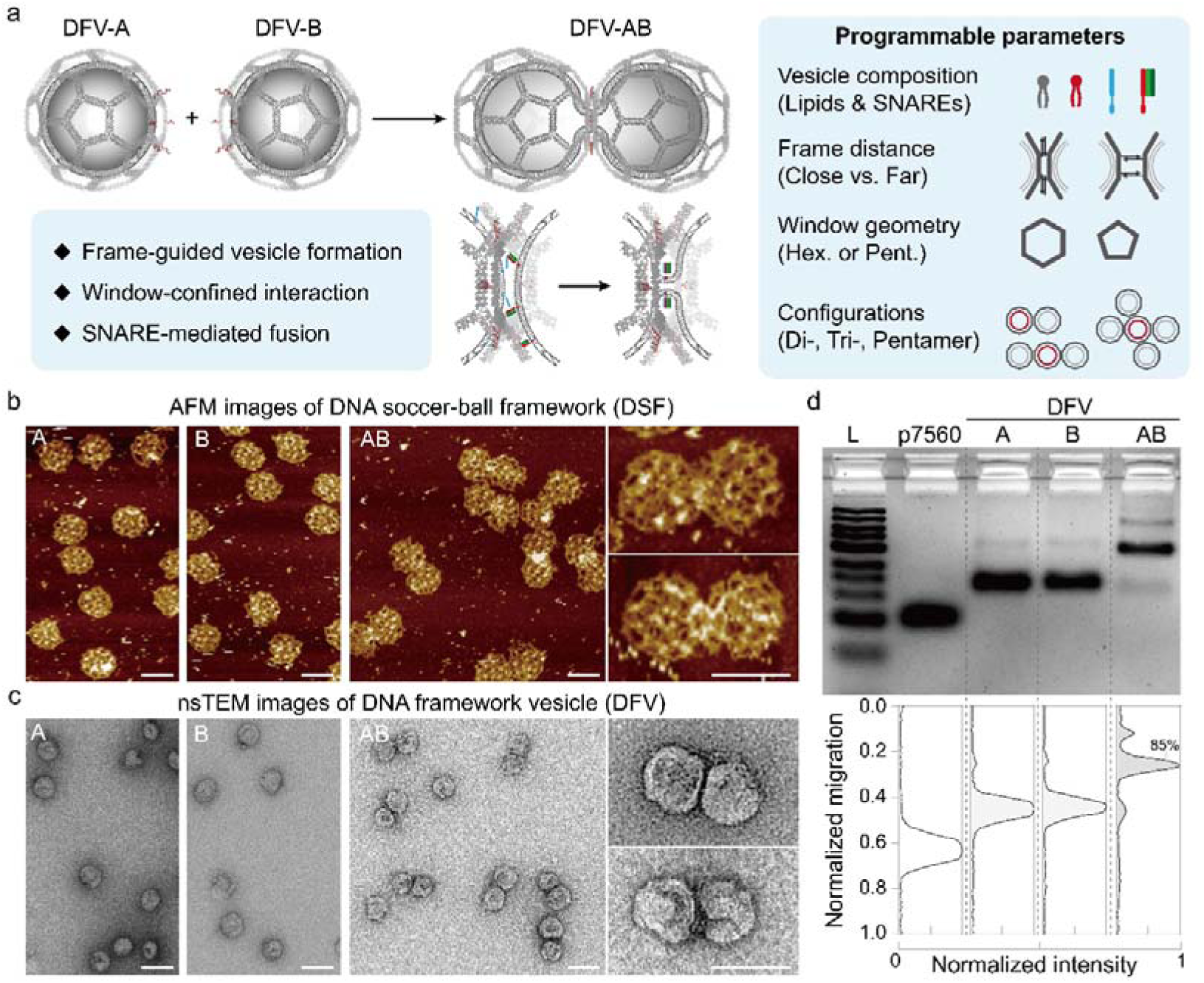
Schematic overview and assembly of DNA nanoreactor setup. **(a)** Programmable platform design using a DNA soccer-ball framework (DSF) enables confined vesicle formation and window-mediated membrane fusion, with key tunable parameters shown in the right panel. **(b)** AFM images of DSF monomers and dimers formed via close docking of hexagonal windows. Scale bar: 100 nm. **(c)** nsTEM images of DFV monomers and dimers encapsulating uniform liposomes without membrane proteins. Scale bar: 100 nm. **(d)** SDS–agarose gel electrophoresis validating the dimerization efficiency of DFV-AB.

## Results

### Design and Construction of DNA Framework Vesicles (DFVs)

To construct a confined nanoenvironment for membrane fusion, we employed a DNA soccer-ball framework (DSF) with the geometry of a truncated icosahedron. Owing to its isotropic architecture and multiple docking portals — comprising twelve pentagonal and twenty hexagonal windows — the DSF serves as an ideal nano-reactor to mediate vesicle-vesicle interactions. Following a standard assembly protocol (*14*), the DSF was constructed using a 7,560-nucleotide (nt) circular single-stranded DNA (ssDNA) scaffold folded by 360 staple strands through a 15-hour thermal annealing process. The assembled structure was purified via rate-zonal ultracentrifugation on a glycerol gradient (see **Methods** in **Supporting Information**, **SI**) and verified by atomic force microscopy (AFM) and negatively stained transmission electron microscopy (nsTEM), as illustrated in **Figure 1b** and **Figure S1**.

Vesicle formation within the DSF cavity was directed by hybridizing cholesterol-modified antisense oligonucleotides (21 nt) to designated inner-handles along the DNA edges, which acted as membrane nucleation seeds (**Table S1**). The seeded DSFs were mixed with octyl-β-D-glucoside (OG) and a lipid mixture consisting of 78% DOPC, 10% DOPS, 10% DOPE, and 2% PEG2k-DOPE (**Tables S2**, **S3**). Detergent removal by dialysis induced membrane assembly inside the DSF cavity, resulting in DNA Framework Vesicles (DFVs). The number and spatial distribution of cholesterol seeds (n= 24) were optimized to support uniform nucleation and growth, yielding highly monodisperse liposomes consistent with previous reports (*15*) (**Figure 1c**). As shown in **Figure S2**, well assembled DFVs products were purified from the untemplated liposomes via density gradient ultracentrifugation against iodiaxanol medium, followed by a fractionation and visualization via SDS-agarose gel electrophoresis (AGE).

To facilitate vesicle docking, single-stranded DNA outer handles were extended from the edges of selected windows on the DSF (**Figure S3**), creating programmable portals for the hybridization-mediated connection of DFVs bearing complementary docking strands (e.g., x/x’, y/y’ or f/f’, sequences in **Table S1**). After 18-hours incubation at 37°C with a 1:1 molar ratio of each counterpart, bare DSF-AB (without inner vesicles) were visualized by AFM (**Figure 1b**), while DFV-dimers (without SNAREs) were examined by nsTEM (**Figure 1c**). Both configurations exhibited high-quality products that efficiently assembled through window docking. The DFV-AB dimers achieved a yield of approximately 85%, as quantified by SDS–AGE band intensity analysis (**Figure 1d**).

### Structural Visualization of Membrane Fusion Intermediates

Building on this design, we next reconstituted the canonical SNARE proteins into DFVs to enable biologically driven membrane fusion and visualize intermediate stages by cryogenic electron microscopy (cryo-EM). As previously established, vesicle-associated membrane protein 2 (VAMP2) on vesicle membranes and a complex of Syntaxin-1 and SNAP-25 (t-SNAREs) on target membranes cooperatively assemble into a four-helix bundle that drives membrane merger through progressive zippering (*5, 16*). In our system, functional SNARE proteins were co-reconstituted into DFVs during the detergent-mediated lipid templating process. Specifically, VAMP2 and t-SNAREs were introduced into DFV-A and DFV-B at protein-to-lipid ratios of 1:200 and 1:400 (*8, 17*), respectively (**Figure 2a**). Proteo-DFVs showed slightly higher buoyant density compared to bare DFVs, with peak fractions appearing at F9–10 (**Figure S4**) versus F8–9 (**Figure S2**) in iodixanol gradient ultracentrifugation. Electron microscopy confirmed their structural integrity and vesicle uniformity, with standard deviation below 7 nm for both DFV-A*_VAMP2_* and DFV-B*_tSNAREs_* (**Figure S5**). Notably, the diameters measured from nsTEM (62–66□nm) reflect the typical flattening of liposomes during staining and drying, which artificially enlarges their apparent size by 1.4-1.5-fold compared to their native dimensions under cryogenic conditions (*10*). Silver staining quantification revealed approximately 33 copies of VAMP2 per DFV-A and 19 copies of t-SNARE per DFV-B, confirming successful reconstitution.

**Figure 2.**
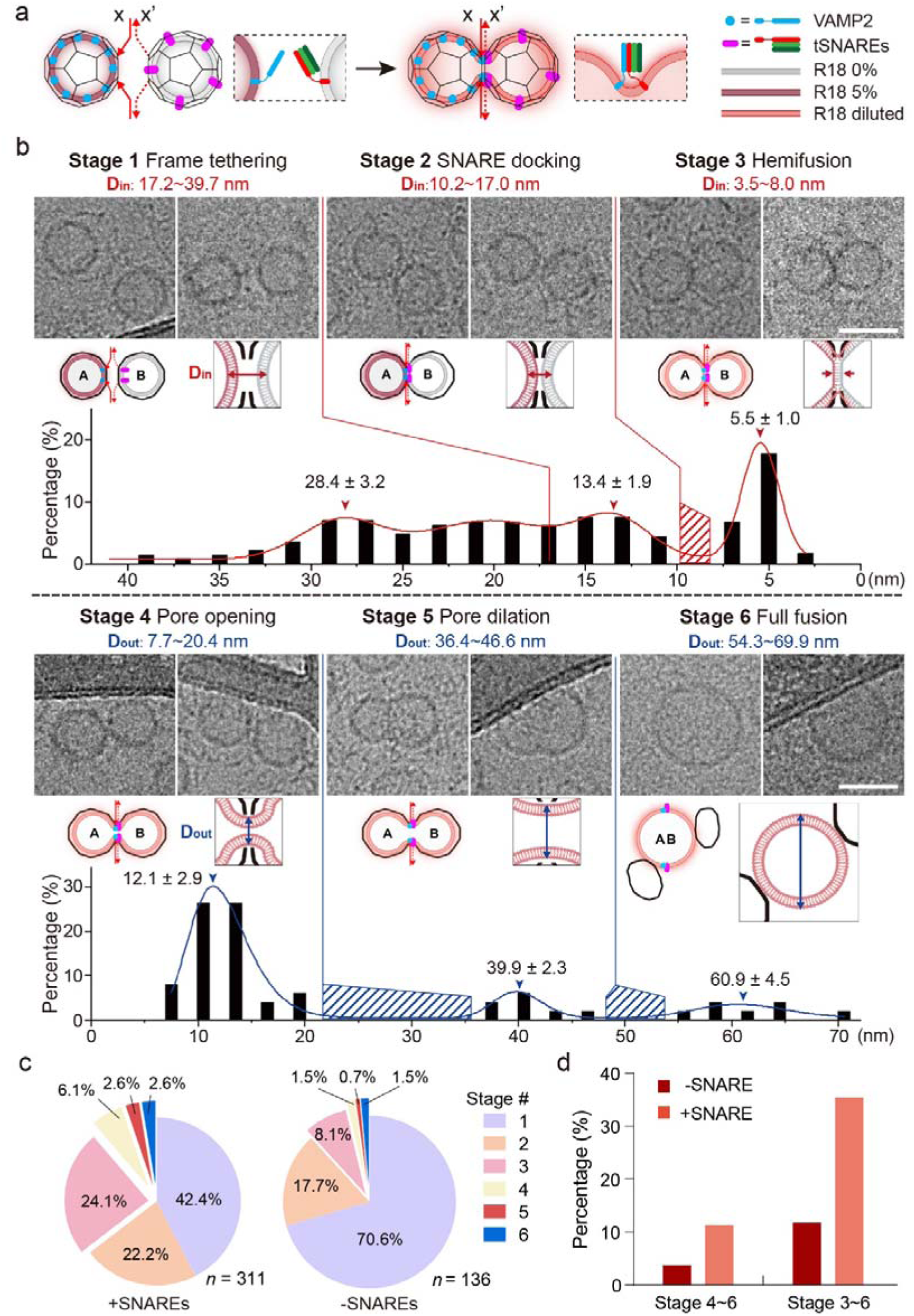
Structural resolution and statistical distribution of SNARE-mediated vesicle fusion intermediates within DNA framework nanoreactors. **(a)** Schematic of DFV-AB fusion with incorporated SNARE proteins and R18 membrane dye. **(b)** Representative cryo-EM images (top), schematic diagrams (middle), and quantified **D_in_** and **D_out_** histograms (bottom) for six structurally resolved fusion intermediates. Red and blue lines represent fitted (multi-peak Gaussian) distribution curves for **D_in_** and **D_out_**, with peak positions (mean± SD) annotated. Scale bar: 50 nm. **(c)** Population distributions of fusion stages under +SNARE and –SNARE conditions. **(d)** Comparison of post-fusion populations (Stage 4–6 and extend to Stage 3-6) under SNARE-positive and SNARE-negative conditions.

Inspired by conventional fluorescence-based assays (*18*), we also incorporated R18 (*19*)—a self-quenching lipophilic dye—into our DFVs to track membrane fusion events via dequenching-induced signal recovery, as detailed in later dynamic studies. A schematic overview of our design is illustrated in **Figure 2a**, where color-coded representations of SNARE proteins and vesicle membranes depict the pre- and post-fusion states. These visual cues are also used to provide conceptual guidance for understanding the intermediate states of vesicle fusion.

Experimentally, proteo-DFV-AB products after 18 hours of incubation were examined by cryo-EM. From hundreds of clearly identifiable dimeric particles, we categorized membrane fusion into six structurally distinct stages, capturing a continuum from pre-fusion to complete vesicle merger (**Figure 2b, S6-S8**). Criteria for distinguishing these intermediates relied on two geometric descriptors: **D_in_**, the shortest distance between the inner leaflets of the two vesicles, which is particularly informative in pre-fusion and hemifusion stages; and **D_out_**, which measures the lateral separation between the outer leaflets of the two membranes after fusion. Together, these quantitative parameters enable resolution of fusion intermediates with nanometer-scale precision.

In the initial frame-tethering stage (**Stage 1**, P1= 42.4%,), two DFVs were juxtaposed through hybridization of docking strands at the hexagonal window, with no evident membrane deformation or contact. The average **D_in_** exceeded 17□nm, forming a distribution valley that separated this state from downstream fusion-related intermediates. The dominant peak at 28.4□nm, centered between the minimum and maximum values (17-40 nm), reflected the spatial offset between vesicle centers imposed by the frame geometry. As membranes approached closer in **Stage 2** (SNARE docking, P2= 22.2%), **D_in_** values narrowed to 10.2–17.0□nm with a peak at 13.4□nm. After subtracting ∼5□nm bilayer thickness from each side, the ∼3.4□nm intermembrane gap aligned well with previously reported partial assembly of trans-SNARE complex distances (*20, 21*), suggesting successful SNARE-mediated membrane engagement. In **Stage 3 (**hemifusion**, P3 = 24.1%)**, membranes appeared partially merged: outer leaflets began to fuse while inner leaflets remained distinct. **D_in_** values further narrowed to 3.5–8.0□nm, peaking at 5.5□nm—consistent with the expected thickness of a single lipid bilayer and indicative of a hemifusion diaphragm (*2, 6, 24*). The merged region (L=17.9 ± 4.8 nm; **Figure S6**) indicates a dynamic hemifusion zone stabilized by the DNA framework. The discontinuity between **Stages 2** and **Stage 3** marked a shift in the dominant driving force from SNARE zippering to lipid interfacial tension.

Following hemifusion, the system entered the post-fusion phase, characterized by membrane continuity and the dynamic nucleation and subsequent dilation of fusion pores. In **Stage 4 (**fusion pore opening, P4 = 6.1%), a visible pore bridged the vesicles, with **D_out_** sharply peaking at 12.9□±□2.9□nm—smaller than the ∼24.2□nm inner diameter of the hexagonal window. After accounting for bilayer thickness, the estimated pore diameter (∼0–5.8□nm) matched prior reports of nascent fusion pores (*25–28*), underscoring the DSF’s capacity to stabilize this transient configuration. In **Stage 5** (pore dilation, P5= 2.6%), the pore widened substantially, yielding a dumbbell-shaped vesicle with a waist **D_out_** ranging from 36.4 to 46.6□nm (mean 39.9□±□2.3□nm). This exceeded the outer diameter of the hexagonal docking window (∼28□nm), implying partial tearing of the original docking windows to accommodate membrane expansion. Nonetheless, the remaining DSF framework still exerted structural constraint on the fused vesicle. Finally, in **Stage 6** (full fusion, P6 = 2.6%), a fully merged, spherical vesicle was observed with an average diameter of 60.9□±□4.5□nm—approximately 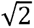-fold of the monomeric DFVs (43□±□3.5□nm, **Figure S5**)—indicative of doubled membrane area. The uniform size of these post-fusion vesicles, resulting from the merger of two identically templated DFVs with conserved lipid content, contrasts with the heterogeneity of free fusion systems. Occasional low-contrast remnants of ruptured DSF scaffolds were attached to the enlarged vesicles. More cryo-EM examples can be found in **Figure S6** and **S7,** and representative 2D class of **Stage 2**, **3**, and **4** particles are summarized in **Figure S8**.

The structural continuum across **Stages 1–6** highlights how geometric confinement eliminates curvature-related variability and directs a well-defined fusion trajectory while stabilizing otherwise transient intermediates. Moreover, the framework itself serves as an energetic reference, enabling delineation of the dominant physical forces governing each intermediate. In **Stage 1**, vesicle tethering is mediated purely by DNA hybridization across complementary handles. **Stage 2** introduces SNARE complex formation, where protein interactions drive membrane approximation. The **Stage 2–3** transition couples SNARE zippering and lipid mixing, producing a hemifusion diaphragm in which membrane tension arising from bilayer deformation is counterbalanced by the rigidity of the framework windows at **Stage 3**. **Stage 3–4** marks a tipping point where residual SNARE zippering, curvature stress, and accumulated tension collectively nucleate the fusion pore. In **Stage 4**, pore size is constrained by DSF geometry and stiffness, indicating that frame rigidity dominates over dilation forces. In **Stage 5**, the expanding pore surpasses the window boundary, and membrane tension prevails, partially tearing the scaffold yet preserving a peanut-like morphology. **Stage 6** represents the terminal, low-population state in which complete membrane merger overrides structural confinement.

As a negative control, cryo-EM analysis of DFV-AB dimers assembled without SNAREs (**Figure S9**) revealed all six morphological stages (**Figure 2c**), yet most vesicle pairs remained in the tethering–approach regime (**Stages 1–2**, P1+P2= 88.3%), reflecting stable frame docking without productive membrane engagement. Under this condition, closer vesicle spacing—formally assigned as “**Stage 2**” in the +SNARE group—represents passive proximity rather than energy-driven docking. Subsequent lipid interaction (**Stage** 3, P3 = 8.1%) occurred only sporadically, while post-fusion products (**Stages 4–6**) accounted for less than 4% of the population—likely arising from random thermal fluctuations or occasional collisions. These results demonstrate that, although confined geometry permits rare spontaneous fusion, efficient and directional merger within the DNA framework strictly depends on SNARE-mediated interactions. The sharp contrast between +SNARE and –SNARE conditions (**Figure 2d**) underscores the driving role of SNARE complexes in overcoming energetic barriers, establishing the DFV nanoreactor as a robust, tunable platform for quantitative fusion analysis under defined geometric and biochemical constraints.

### Kinetic Profiling of DFV-Directed Membrane Fusion

Beyond single-particle visualization by cryo-EM, bulk fluorescence assays provide complementary insights into the kinetics and probability of framework-directed membrane fusion. Fluorescence dequenching or Förster resonance energy transfer (FRET), driven by lipid mobility and inter-molecular distance changes, are classical strategies to monitor fusion events. In our preliminary evaluation, the R18 (*19*) dequenching assay yielded a substantially higher signal-to-noise ratio and resolution under the current lipid composition and reaction conditions than the NBD/Rhodamine (*18*) FRET assay (**Figure S10**). We therefore adopted the R18-based system for subsequent fusion kinetic measurements.

Following the standard approach, we mixed R18-labeled lipids with unlabeled lipids at defined molar ratios (1:0, 1:0.5, 1:1, 1:2, 1:4, 1:8) to establish a correlation between fluorescence intensity and R18 dilution, thereby defining the quantitative relationship between fluorescence recovery and the *rounds of fusion* (ROF) (**Figure S11**). When one DFV partner contained R18-labeled lipids, real-time fluorescence tracing of proteo-DFV-AB nanoreactors revealed both the kinetic evolution and endpoint fluorescence values, which were converted to ROF using the established standard curve. In conventional bulk liposome fusion assays, ROF values represent ensemble-averaged fluorescence changes, where the identities of vesicles and occurrence of multiple fusion rounds on a single vesicle remain indistinguishable. In contrast, the geometrically confined DFV-AB system enables direct interpretation: the ROF percentage effectively indicates how many DFV pairs (per 100) underwent successful fusion, providing a more precise and quantitative metric for comparative analysis.

Taking advantage of the structural editability of DNA origami frameworks, we next investigated how key geometric parameters—including inter-frame distance, docking window configuration, and multivalent assembly—quantitatively affect fusion kinetics and efficiency (**Figure 3a**). As suggested by the cryo-EM analyses in the previous section, pre-docking of frameworks plays a crucial role in promoting membrane fusion. We therefore first examined the effect of increasing inter-vesicle distance on fusion probability. Using DFV*_H/C_*-AB assembled via six close-type linkers (3′-x / 5′-x′) at the hexagonal window as a reference, we constructed a far-type variant, DFV*_H/F_*-AB, where the linker pair (3′-f / 3′-f′, **Figure S12**) extended the spacing between opposing frameworks from ∼2 nm (the diameter of a dsDNA duplex) to ∼7 nm (the length of a 21-bp dsDNA segment). A third control, DFV*_NL_*-AB, lacked inter-frame linkers altogether, allowing only random encounters between vesicles through arbitrary windows.

**Figure 3.**
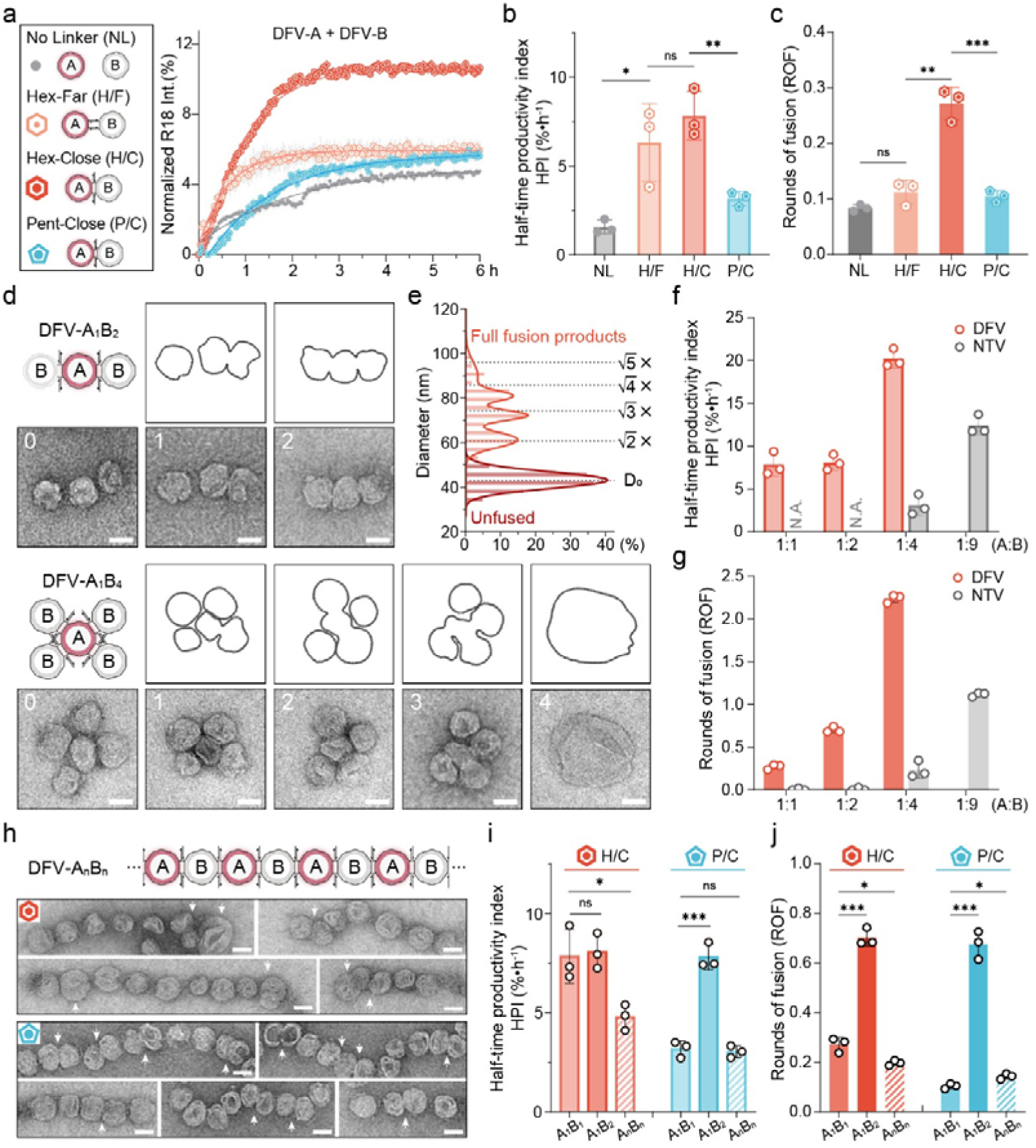
Framework geometry governs membrane-fusion dynamics. **(a)** Schematics of DFVs with varied inter-frame distances and window geometries, and the corresponding R18 fluorescence traces. **(b)** Comparison of kinetic profiles using the half-time productivity index (HPI). **(c)** Quantified rounds-of-fusion (ROF) derived from endpoint fluorescence measurements. **(d)** nsTEM images of multivalent DFV assemblies formed under different fusion combinations. Scale bar: 50 nm. **(e)** Diameter distributions obtained from cryo-EM measurements of inflated fusion vesicles. **(f–g)** Fusion behavior of DFVs with different stoichiometries compared with free-vesicle mixtures. **(h–j)** Representative images and quantitative analysis of linearly assembled DFV arrays via hexagonal and pentagonal docking windows. Scale bar: 50 nm.

Real-time fluorescence tracing of R18 dequenching was performed using a microplate reader. As shown in **Figure 3a**, all three systems displayed similar initial fluorescence increase within the first 30 minutes, suggesting comparable early-stage docking and lipid rearrangement. However, DFV*_H/F_*-AB exhibited a rapid attenuation in fluorescence growth thereafter, reaching a markedly lower final intensity than the closely tethered DFV*_H/C_*-AB, indicating reduced overall fusion probability when the membranes were held further apart. The no-linker DFV*_NL_*-AB sample displayed the weakest fusion signal, closely overlapping with the SNARE-deficient DFV*_H/C_*-AB*_bare_* control (**Figure S13**). These results underscore the cooperative contributions of both frame tethering and SNARE docking in driving membrane merger—consistent with the sequential membrane-approach events captured in **Stage 1** and **Stage 2**.

In analyzing the overall reaction kinetics, we found that the initial rate, rate variation, and final fluorescence intensity differed among the tested conditions, rendering a single-exponential rate constant (*k*) insufficient for comprehensive comparison of fusion efficiency. To integrate both the reaction speed and magnitude of fluorescence change, we introduced a *half-time productivity index* (HPI), defined as 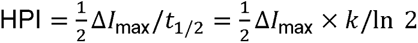 (**Figure 3b**). Although the differences between groups did not reach statistical significance (*p > 0.05*), the averaged values followed the trend HPI*_H/C_*> HPI*_H/F_*, with the *close-type* DFV*_H/C_*-AB showing smaller standard deviations across replicates, indicating improved system stability and overall fusion efficiency conferred by near-distance docking.

Using the endpoint fluorescence values and the calibration curve, we further calculated the *rounds of fusion* (ROF) for each condition (**Figure 3c)**. The DFV*_H/C_*-AB group showed an ROF of 0.271, indicating that ∼27% of DFV pairs underwent fusion. This value fell between the cryo-EM–derived proportions of hemifused-to-fused states (P3–P6 = 0.354) and fully fused products (P4–P6 = 0.113) (**Figure 2d**), demonstrating strong consistency between single-particle analysis and bulk fluorescence statistics within the framework-confined system. In comparison, the ROF of DFV*_H/F_*-AB (0.112) indicated that approximately 11.2% of vesicle pairs reached Stage 3–6 (i.e., achieved hemifusion or beyond), while the no-linker DFV*_NL_*-AB exhibited a baseline fusion probability of ROF = 0.084, reflecting random collision events. Although the ROF difference between H/F and NL groups was not statistically significant (*p > 0.05*), the higher HPI observed for DFV*_H/F_*-AB, together with its characteristic kinetic curve, suggests that distant tethering promotes early-stage collisions but may hinder subsequent membrane merger—revealing a dual effect of long-range linkage on fusion dynamics.

We next examined how docking window geometry influences fusion efficiency under the *close-type* configuration. By pairing two pentagonal windows to assemble DFV*_P/C_*-AB dimers (see details in **Figure S14**), we observed a marked reduction in both fusion kinetics and overall productivity compared with the hexagonal-window DFV*_H/F_*-AB. Cryo-EM inspection of DFV*_P/C_*-AB revealed that the *Stage 4* products exhibited nearly identical D_out_ values to those of the hexagonal group (12.8 vs. 12.6 nm, **Figure S15**). Considering that the inscribed diameters of pentagonal and hexagonal windows (19.3 nm and 24.2 nm, respectively) both exceed the measured fusion pore size, the SNARE-dependent pore formation thus appears unaffected by the outer window dimension.

In contrast, the reaction kinetics indicated a substantially slower initial rate in the pentagonal system, attributable to two geometric and stoichiometric constraints. First, the number of inter-frame linkers decreased from six to five, reducing the dimerization efficiency from ∼85% to ∼69% (AGE analysis, **Figure S17**). Second, the facing membrane area defined by the docking window was smaller (A*_pent_* = 293 nm² vs. A*_hex_* = 462 nm²), lowering the probability of v-/t-SNARE pairing within the confined zone by approximately 37%. Together, these effects resulted in a pronounced decrease in overall fusion efficiency, reflected by significantly reduced HPI. Correspondingly, the measured ROF of DFV*_P/C_*-AB (0.104) was lower than that of DFV*_H/C_*-AB (0.271).

To further assess the capacity of the DFV platform for multi-round fusion, we introduced multi-window DFV-A designs (R18-labeled) under the *close-type* configuration, constructing DFV-A_1_B_2_ trimers and DFV-A_1_B_4_ pentamers by pairing opposite or mixed hexagonal/pentagonal windows (**Figure S16**).Agarose gel electrophoresis (AGE) analysis (**Figure S17**) revealed that DFV*_H/C_*-A_1_B_2_ assembled efficiently, achieving ∼80% yield comparable to the dimer, whereas the pentagonal counterpart DFV*_P/C_*-A_1_B_2_ produced less than 50% complete products. The DFV-A_1_B_4_ pentamer exhibited even lower yield (<18%), with substantial accumulation of lower-order species, particularly tetramers (A_1_B_3_), reaffirming the superior docking efficiency of hexagonal windows.

Despite lower yields, nsTEM imaging resolved distinct fusion intermediates corresponding to 0–2 fusion events in A_1_B_2_ and up to four in A_1_B_4_ (**Figures 3d**). Cryo-EM characterization further confirmed the heterogeneous yet well-defined fusion combinations. For DFV-A_1_B_2_ (**Figure S18**), the asynchronous interactions between the central A-vesicle and its two B-partners led to diverse combinations of stages (S1–S6) on each side, all of which could be individually resolved. For DFV-A_1_B_4_ (**Figure S19**), however, the increased structural complexity and off-plane deformation induced by multi-point fusion precluded a systematic stage classification; therefore, only representative morphologies are shown. Statistical analysis of unfused vesicles (**Stages 1–2**) and fully fused vesicles (**Stage 6**) from both A_1_B_2_ and A_1_B_4_ systems (**Figure 3e, S20**) revealed that unfused vesicles retained the same diameter distribution as DFV monomers, peaking at D₀ = 43 nm. In contrast, the diameters of fully fused vesicles exhibited four Gaussian peaks corresponding closely to the theoretical sizes expected from one to four sequential fusion events, following the relation D ≈ √(n + 1) × D₀, (n = 1–4). These results highlight the geometric precision and quantitative predictability of the DFV nanoreactor platform in mediating integer-step, vesicle-by-vesicle membrane fusion.

Bulk fluorescence tracing of R18 dequenching was performed for DFV-A1B1, DFV-A_1_B_2_ (Hex), and DFV-A_1_B_4_ assemblies, and compared with non-templated vesicle (NTV) systems in which freely formed v-SNARE/R18–labeled vesicles (A) and t-SNARE vesicles (B) were mixed at molar ratios of 1:1, 1:2, 1:4, and 1:9. As shown in **Figures 3f and S21**, fusion efficiency in DFV nanoreactors increased markedly with higher B-to-A ratios and was consistently much greater than that of the free vesicle system. This enhancement arises from two key factors: (i) the presence of multiple, directionally aligned docking windows that increase the probability of membrane encounters, and (ii) within higher-order DFV assemblies, each newly fused product remains spatially proximate to the remaining unfused B-vesicles, whereas in the free system, each fusion event consumes reactants and lowers the effective particle concentration, thus reducing subsequent fusion likelihood. Interpreted through the single-particle statistics enabled by the DFV framework, the ROF values of 0.70 and 2.24 for DFV-A_1_B_2_ (Hex) and DFV-A_1_B_4_ reflect increasing fusion multiplicity per A-vesicle (**Figure 3g**). This integer-based interpretation of ROF is made possible by the uniformity and quantized assembly of the DFV platform, providing an intuitive and statistically meaningful measure of fusion probability. In contrast, ROF values in conventional NTV assays reflect only ensemble-averaged fluorescence changes, lacking direct correspondence to discrete fusion events—highlighting the unique analytical precision and mechanistic transparency enabled by the DFV nanoreactor system.

In contrast to the finite A_1_B_n_ assemblies, we further investigated membrane fusion in linearly extended DFV arrays (A_n_B_n_) formed through opposite-window docking (**Figure S22**). Such linearly extended assemblies resemble the sequential compound fusion observed in excitable cells, where consecutive vesicle–vesicle fusion events at pre-fused membranes enhance the release capacity and dynamic range of exocytosis (*29*). nsTEM imaging revealed long chain–like assemblies under both hexagonal and pentagonal docking conditions, with the hexagonal DFV*_H/C_* - A_n_B_n_ structures exhibiting visibly greater lengths (see AGE results and length distribution in **Figure S23**). As illustrated by representative images in **Figure 3h**, hemifused and fused vesicles (**Stages 3–6**, indicated by white arrows) could be clearly identified along the linear assemblies, where adjacent vesicles served as mutual references.

Compared with the trimeric DFV- A_1_B_2_, however, the linear DFV- A_n_B_n_ arrays showed unexpectedly lower overall fusion efficiency (**Figure 3i and S24**): their composite HPI values were below those of the A_1_B_1_ dimers and substantially below the A_1_B_2_ systems for both hexagonal and pentagonal configurations (**Figure 3j**). Several factors likely contribute to this decline. First, the A_1_B_2_ assemblies inherently benefit from a twofold excess of B-vesicles, providing greater dye-dilution capacity than the stoichiometric A_n_B_n_ arrays. Second, under identical total DFV-A and DFV-B concentrations, the formation of large A_n_B_n_ polymers reduces the number of individual complexes, thereby lowering the fraction of random, non-templated collisions. Third, within a long linear chain, the close approach of one A–B pair (**Stage 2**) necessarily increases the spacing of its neighboring vesicles, diminishing their subsequent contact probability and occasionally producing isolated vesicles with both sides in a distant configuration—an effect evident in **Figure 3h**. ROF analysis further supported these observations: in the hexagonal series, the A_n_B_n_ assemblies displayed reduced fusion probability relative to A_1_B_1_ and A_1_B_2_, whereas in the pentagonal system, the shorter chain length and slower reaction kinetics preserved a slightly higher chance for repeated fusion, yielding marginally higher ROF values to its A_1_B_1_ counterparts.

Together, these systematic kinetic and structural analyses establish a clear quantitative framework for understanding how DNA-origami–defined geometry governs membrane fusion efficiency. The confined DFV system reveals that both the proximity and multiplicity of docking windows critically determine fusion probability: near-distance (close-type) linkers and hexagonal interfaces consistently outperform distant or pentagonal configurations. Multivalent and higher-order assemblies enable multiple fusion events but introduce geometric and kinetic constraints that can either amplify or suppress overall productivity, depending on stoichiometry and spatial arrangement.

### Population-Level Quantification of Fusion Events by Nano-Flow Cytometry

Moving beyond ensemble fluorescence assays and low-throughput EM imaging, we employed nano-flow cytometry (*30, 31*) (nFCM) to achieve single-vesicle, high-throughput quantification of membrane fusion in DNA nanoreactors with varied stoichiometries (**Figure 4a**). Instead of using R18 lipid dequenching, which yields broad and continuously distributed fluorescence intensities after fusion, we independently labeled DFV-A*_VAMP2_* and DFV-B*_t-SNARE_* with FITC and Cy5 modified cholesterol, two spectrally separated fluorophores that do not interfere with each other. This dual-color strategy enables precise identification of fused products based on fluorescence co-existence (FITC⁺Cy5⁺) rather than intensity changes, providing a direct and quantitative readout of fusion events.

**Figure 4.**
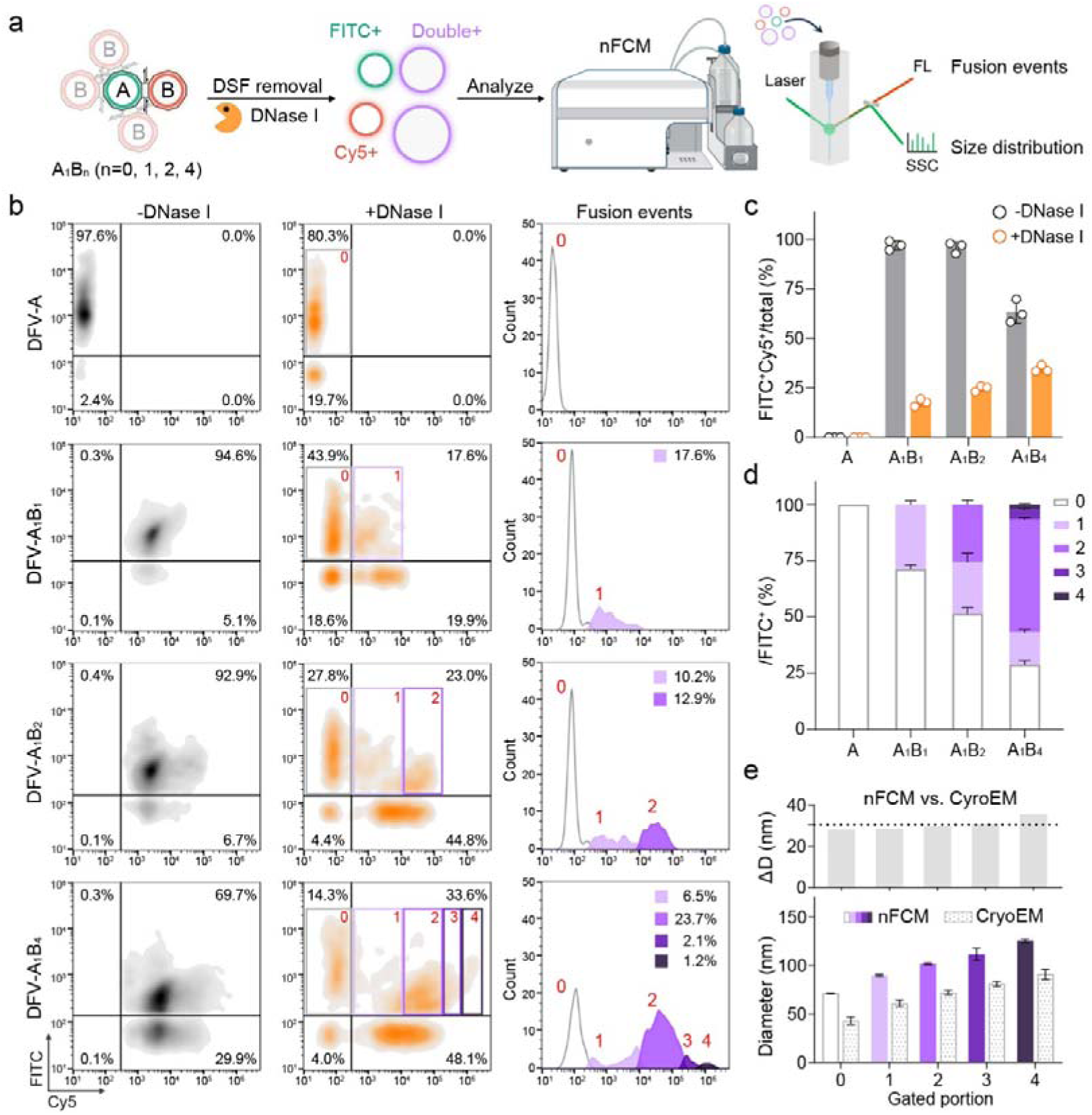
Single-vesicle analysis of DFV-mediated membrane fusion by nano-flow cytometry. **(a)** Dual-color design of DFVs labeled with FITC and Cy5. **(b)** Representative fluorescence-intensity plots before and after DNase I digestion. **(c)** Distribution of fluorescence populations at varied A:B ratios. **(d)** Fractional analysis of 0–4 fusion products within the FITC-positive population. **(e)** Size profiles of vesicles corresponding to different fusion rounds.

As shown in **Figure 4b** (left column) and quantified in **Figure 4c**, monomeric DFV-A vesicles exhibited only FITC⁺ signals, whereas co-assembled DFV-A/B nanoreactors displayed an additional double-positive population (FITC⁺Cy5⁺) indicative of successful fusion. At A:B ratios of 1:1 and 1:2, over 90 % of particles formed nanoreactor assemblies, while further increasing the B ratio to 1:4 led to a pronounced reduction of residual B-rich super-assemblies. This population trend closely matched the distribution of dimeric and multimeric bands observed in SDS-AGE analysis (**Figure S17**), confirming that the nFCM profiles faithfully reflect the hierarchical assembly states of DFV systems.

Upon DNase I digestion (**Figure S25**), the DNA frameworks were removed, leading to the disassembly of higher-order superstructures and separation of individual vesicles. Under these conditions, the double-positive (FITC⁺Cy5⁺) population corresponds to vesicles that underwent actual lipid merger (**Stages 3–6**), representing genuine membrane fusion events (**Figure 4b**, second column). For comparison, monomeric DFV-A controls exhibited nearly identical fluorescence distributions before and after DNase I treatment, with only a minor population (19.7%) appearing in the double-negative quadrant (Q3), likely attributable to small enzyme–nucleic acid aggregates detected by the highly sensitive nFCM system. In contrast, the DFV-A_1_B_1_ dimer population resolved clearly into three subgroups—FITC⁺ (Q2), Cy5⁺ (Q4), and FITC⁺Cy5⁺ (Q1)—after framework removal. Interestingly, the FITC⁺ subpopulation displayed a slight rightward shift along the Cy5 axis compared with the DFV-A control, which likely arises from limited cholesterol exchange inherent to the vesicle staining strategy, rather than representing genuine membrane fusion.

Within the double-positive gate, the proportion of fused vesicles increased progressively with higher B-to-A ratios (**Figure 4c**), consistent with the ROF trend observed in bulk R18 assays (**Figure 3g**). Notably, as the number of B vesicles increased, the Cy5 fluorescence of double-positive populations exhibited a stepwise rightward shift along the Cy5 axis. Each higher-order assembly produced additional Cy5⁺ subpopulations that appeared precisely beyond the Cy5 range of the preceding, lower-order group, confirming that newly emerged Q1 signals correspond to genuine fusion products rather than spectral overlap.

For the A_1_B_1_ system, which theoretically allows only a single fusion event, its Cy5 intensity window served as a reference for gating. In the A_1_B_2_ sample, an additional Cy5-enriched subpopulation appeared just beyond this range, differing by approximately one order of magnitude in Cy5 intensity and representing vesicles that underwent a second fusion event (**Figure 4b**, third column). The A_1_B_4_ sample further exhibited a distinct bimodal distribution, introducing two new populations that together spanned two additional orders of Cy5 signal. These features strongly suggest that the two peaks correspond to vesicles completing three and four rounds of fusion, respectively.

Importantly, the four consecutive Cy5 intensity intervals are evenly spaced by roughly one order of magnitude, indicating clear signal separability among 1-, 2-, 3-, and 4-fusion products. The regular 10× intensity increments likely reflect the inherent signal gaining settings of the nFCM detector rather than proportional increases in absolute fluorescence yield, yet this systematic scaling demonstrates strong internal consistency and confirms the reliability of the gating strategy for precise, single-vesicle quantification of integer-step fusion.

Because the input amount of DFV-A was kept constant across all samples, we were able to analyze the relative proportions of 0–4 fusion products within the FITC⁺ population (**Figure 4d**). Remarkably, the A_1_B_1_ group showed a single-fusion fraction of 27 %, nearly identical to the previously determined ROF = 0.27, underscoring the strong internal consistency between nFCM and bulk fluorescence analyses. The A_1_B_2_ group likewise exhibited a quantitatively consistent distribution, where one- and two-fusion products together accounted for roughly 70 % of total FITC⁺ vesicles—well aligned with its ROF-derived probability. Although the A_1_B_4_ system displayed slight deviations in absolute values, the dominance of two-fusion products remained in clear agreement with its high overall ROF (> 2). Collectively, these results reinforce the reliability of the nFCM-based population analysis and its congruence with bulk kinetic measurements.

Leveraging the size-resolving capability of nFCM, we further analyzed the dimensional evolution of vesicles across different fusion subpopulations. A calibration curve was first established using S-16M-Exo silica nanosphere standards (**Figure S26**), followed by size analysis of the subgroups corresponding to distinct fusion rounds (**Figure S27**). As shown in **Figure 4e** (bottom), the median diameter increased systematically with the number of fusion events: ∼71.4 nm for unfused monomers, ∼89.8 nm after one fusion, ∼101.9 nm after two, and further to ∼111.6 nm and ∼125.6 nm for three and four fusion events, respectively. This stepwise increase closely matched the cryo-EM–derived dimensions (data summarized from Figure 3e; Gaussian peak means ± SD), yielding an average deviation of ΔD = 30.4 ± 3 nm (**Figure 4e**, **top**). Because nFCM size estimation relies on light-scattering intensity relative to silica standards, this offset can be attributed to systematic differences between vesicle and silica refractive properties as well as instrument calibration. Given the geometric accuracy of cryo-EM measurements, these data suggest that a constant offset of ∼30 nm may be used as an empirical correction factor for nFCM vesicle sizing.

Collectively, dual fluorescence nFCM analysis not only validated membrane fusion within DNA nanoreactors but also quantitatively resolved fusion multiplicity and product size, highlights the platform’s remarkable precision in programming and controlling membrane behavior. When integrated with R18-based kinetic assays and high-resolution cryo-EM imaging, this high-throughput, single-particle approach provides an unprecedented quantitative perspective for dissecting the dynamics and mechanisms of membrane fusion in the DFV system.

### Cell-Surface Membrane Fusion Enables Functional siRNA Delivery

Inspired by natural enveloped viruses and neuronal vesicles that initiate precise membrane fusion at the cell surface to deliver macromolecular cargos, we next explored whether our precisely assembled DFV nanoreactors could function as an artificial membrane fusion system targeting living cells. To achieve this, we established a HeLa cell model stably expressing the membrane-anchored fusion protein EGFP-VAMP2 (*32*) (**Figure S28**). By transfecting a plasmid encoding an N-terminally EGFP-tagged “flipped” VAMP2, the v-SNARE component was oriented outward on the plasma membrane, providing a defined docking interface for DFVs bearing complementary t-SNAREs. In parallel, cholesterol-modified DNA linkers (x′ or f′) were selectively inserted into the cell surface to mediate controllable DFV–cell distances (*33*), designated as *close* or *far* configurations, while cells treated with non-complementary random sequences served as controls.

To specifically observe membrane fusion events occurring on the cell surface rather than those arising from endocytosis, we first screened multiple endocytosis inhibitors (*34*) (**Figure S29**) and selected cytochalasin D (Cyto D), which showed strong inhibitory efficacy with minimal cytotoxicity, for pretreatment. Subsequently, DFV-A_t-SNARE_ vesicles carrying 5% R18 (0.5 nM, 100 µL) were incubated with the cells to visualize R18 dequenching caused by membrane fusion and dye dilution at the plasma membrane (**Figure 5a**). Confocal imaging and mean fluorescence intensity (MFI) quantification revealed that in HeLa-VAMP2 cells, DFV-A_t-SNARE_ tethered in the *close* configuration (x/x′ pair, ∼2 nm distance) produced a strong, continuous R18 signal outlining the cell membrane, indicating efficient membrane fusion.

**Figure 5.**
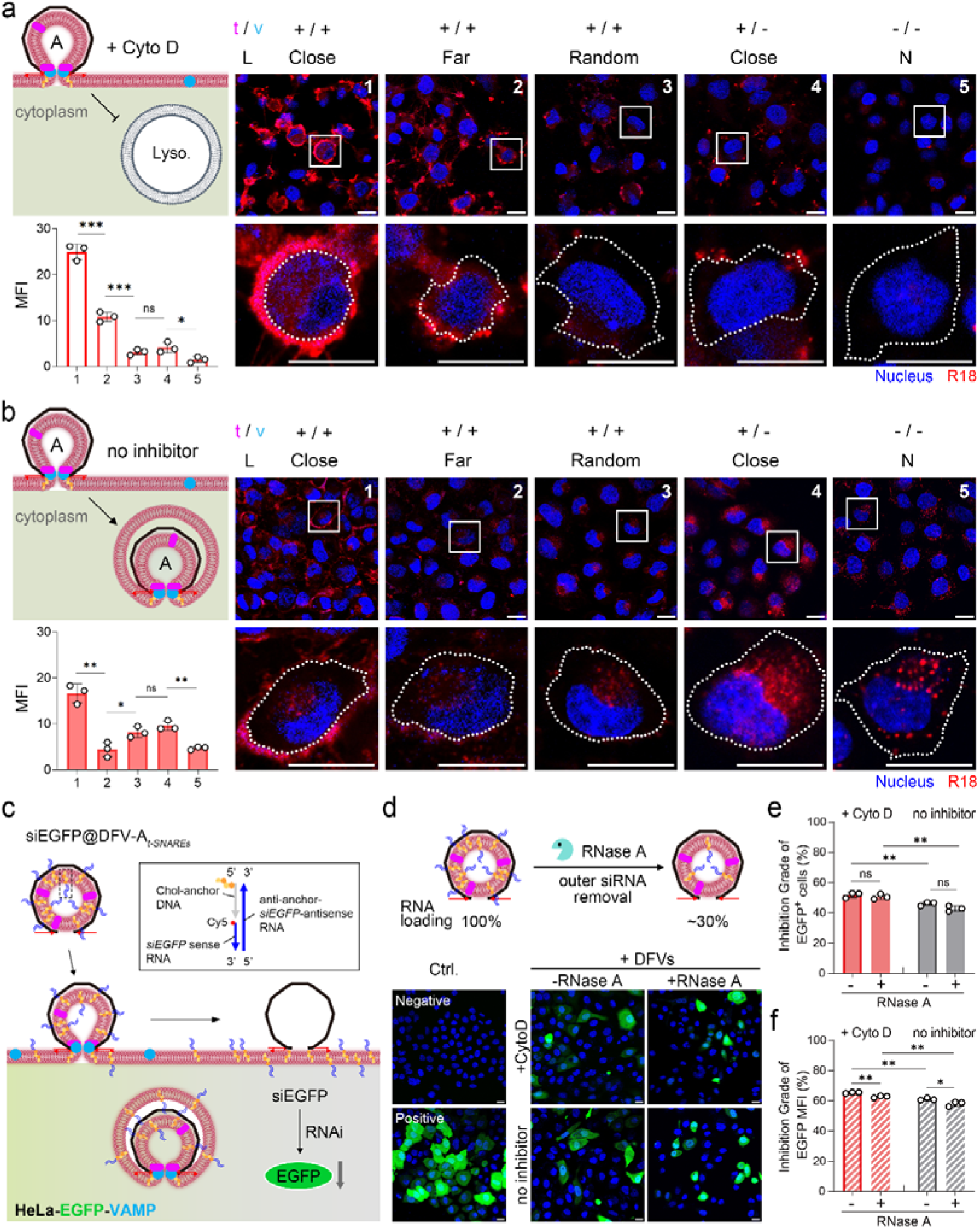
DFV-induced membrane fusion and siRNA transfer in living cells. (a-b) Confocal images of R18-labeled DFV–cell interactions under different tethering conditions with or without Cyto D-mediated endocytosis inhibition. Scale bar: 20 µm. **(c)** Schematic illustration of siRNA incorporation into DFV-A*_t-SNAREs_* via cholesterol-bridged linkage. **(d)** Schematic illustration with quantitative analysis of RNase A digestion targeting siRNA located on the outer surface of DFVs; confocal fluorescence images of HeLa-EGFP-VAMP2 cells treated with siRNA-loaded DFVs under ±RNase A and ±Cyto D conditions. Scale bar: 20 µm. **(e–f)** Flow-cytometric measurements of EGFP expression and mean fluorescence intensity in treated cells.

In contrast, when DFVs were linked to the cell surface via the *far* configuration (f/f′ pair, ∼7 nm distance), the membrane fluorescence intensity markedly decreased—showing a 2.3-fold reduction in MFI—highlighting the decisive role of nanometer-scale proximity in initiating fusion. Cells modified with non-complementary random sequences (*+/+, Random*) or expressing no VAMP2 but tethered in the *close* mode (*+/–, Close*) displayed only punctate fluorescence signals with an approximately 8-fold lower MFI than the *+/+, Close* group, reflecting limited and stochastic fusion driven solely by nonspecific SNARE interaction or DNA proximity. In the double-negative control (*–/–, N*), virtually no membrane fluorescence was observed, excluding nonspecific adhesion or background staining. Together, this controllable fusion provides a conceptual route for membrane augmentation—enabling the transfer of exogenous phospholipid components or membrane proteins to reshape cellular interfaces, offering a promising strategy for programmable cell-surface modification.

To better simulate general cell-engineering and delivery conditions where endocytosis is not inhibited, we next examined DFV-induced membrane fusion in the absence of Cyto D. Under the *close* configuration, in addition to the strong plasma membrane fluorescence, abundant punctate intracellular R18 signals were observed, indicating that part of the DFV population underwent endocytosis. Notably, as illustrated in **Figure 5b**, some DFVs underwent endocytosis shortly after membrane tethering. In these cases, if the invaginating endocytic membrane happened to carry surface-expressed VAMP2 molecules, membrane fusion could subsequently occur at the endosome–lysosome stage, as schematically depicted. Compared with Cyto D-treated cells, the overall MFI decreased from ∼25% to ∼15%, which can be attributed to a higher proportion of internalized DFVs and the limited membrane area of endocytic vesicles, resulting in reduced R18 dilution capacity relative to the plasma membrane. A parallel experiment with sucrose pretreatment to inhibit clathrin-mediated endocytosis yielded similar fluorescence patterns and quantitative trends (**Figure S30**).

In the *far* and *random* tethering configurations, membrane fluorescence was nearly undetectable, suggesting that, in the kinetic competition between fusion and endocytosis, only *close* tethering retained sufficient pre-endocytic proximity for plasma membrane fusion, whereas the rapid onset of endocytosis precluded observable surface fusion once the vesicle–cell distance increased. The even weaker intracellular fluorescence in the *far* group further highlights the inhibitory effect of increased spacing on fusion probability. In VAMP2-negative cells, intracellular fluorescence mainly resulted from lysosomal membrane degradation and phospholipid redistribution, rather than SNARE-dependent fusion; the slightly higher R18 signal in the *close* group reflected enhanced uptake efficiency conferred by DNA tethering.

Taken together, these results demonstrate that nucleic acid–mediated tethering of DFVs in close proximity significantly enhances protein-dependent membrane fusion efficiency. Inspired by these findings, we next adapted the DFV platform for functional cargo delivery using siRNA as a model. An anti-EGFP siRNA (siEGFP) targeting the co-expressed EGFP/VAMP2 cell line was designed and its silencing efficiency verified with Lipofectamine 2000 (**Figure S32**). We then constructed a DFV-based siRNA loading and delivery system to evaluate its fusion-driven gene knockdown capability.

In this design, siEGFP was anchored to a cholesterol-modified DNA linker via a tri-strand bridging strategy and subsequently incorporated into DFV-At-SNAREs during detergent removal (**Figure 5c**). Co-migration of Cy5-labeled siEGFP with phospholipids in ultracentrifugation fractions (**Figure S31**) verified efficient incorporation of siRNA to the DFV lipid membrane. RNase A digestion could markedly reduce the Cy5 signal, revealing that ∼30% of siRNA remained protected within the vesicle lumen (**Figure 5d**).

HeLa-EGFP-VAMP cells were then incubated with DFV samples carrying a total of 2 pmol siEGFP, with or without Cyto D pretreatment to inhibit endocytosis. Flow-cytometric analysis revealed that DFV-A*_t-SNAREs_* achieved pronounced EGFP knockdown under endocytosis-blocked conditions, reducing the proportion of EGFP-positive cells by 51.9%. Remarkably, this silencing efficiency remained nearly unchanged after RNase A pretreatment, despite the effective siRNA dose decreasing to ∼0.6 pmol (**Figure 5e**). Consistent trends were observed for mean fluorescence intensity (MFI), with only ∼1% difference between RNase-treated and untreated samples (**Figure 5f**). These results provide strong evidence that siRNA is directly delivered into the cytosol through membrane fusion rather than via endocytosis and lysosomal escape, and that only siRNA enclosed within DFVs contributes to the functional effect—perfectly mirroring the topology of the fusion process.

Notably, even without endocytosis inhibition, the DFV system maintained robust gene silencing (∼46.2%), outperforming Lipofectamine 2000 under the same siRNA dosage (2 pmol: ∼38%, **Figure S32**). Together with the R18 fluorescence distribution, these results indicate that rapid cell-surface fusion and early endosomal fusion collectively enable efficient cytoplasmic delivery. The modular and programmable nature of the DFV architecture thus offers a versatile platform readily adaptable to other fusion-protein pairs for targeted drug or nucleic acid delivery.

## Discussion

By confining lipid membranes within a nanoscale DNA framework, this study transforms the fleeting act of membrane fusion into a programmable and quantifiable nanoreaction. The DNA framework vesicle (DFV) enables direct visualization and control of a complete fusion sequence—from SNARE-mediated docking to full vesicle merger—capturing intermediates that previously escaped detection. Cryo-EM provides structural snapshots that delineate discrete fusion stages; real-time fluorescence tracing reveals their kinetic transitions; and nano-flow cytometry establishes population-level quantification, together forming a coherent, cross-validated picture of the entire process. The combination of geometric precision and molecular machinery converts stochastic collisions into deterministic, single-pathway events, providing a unified experimental framework that links structural observation with kinetic and statistical evidence.

Beyond structural reconstruction, this work establishes a conceptual framework for geometry-guided membrane reactions, in which the spatial organization of lipid bilayers becomes an active variable in controlling reaction pathways. Recent progress in DNA-origami–templated vesicle assemblies (*35*) has provided elegant architectural control, yet such systems typically isolate membranes from direct lipid continuity. By introducing nanoscale apertures as programmable reaction windows, the DFV nanoreactor transcends this limitation—transforming static vesicle arrays into dynamic, condition-tunable fusion systems. The resulting ability to modulate confinement, curvature, and stoichiometry provides an experimental language to interrogate the energy landscape of bilayer remodeling. Coupled with modern cryogenic imaging, our approach has already resolved intermediate states once accessible only through inference.

Further integration of high-resolution optical (e.g., PAINT or STORM, (*36, 37*)) and cryogenic (*25, 38*) techniques will be invaluable for visualizing the spatial organization and dynamic evolution of both lipids and fusion proteins within confined interfaces. Such multimodal observation could reveal how molecular geometry and mechanical forces cooperate during pore initiation and expansion, deepening our understanding of the physical principles underlying membrane merger. Extending this precision toward real-time structural tracking will transform the DFV platform into a powerful bridge between bottom-up reconstitution and top-down biological observation. In this light, the DFV platform represents a paradigm shift toward programmable membrane biophysics—where fusion events are no longer observed by chance but are designed, stabilized, and quantitatively analyzed through molecular architecture.

In a broader perspective, the DFV platform also demonstrates translational potential as a modular vesicle carrier that delivers cargos through direct membrane fusion rather than endocytic uptake. The ability to achieve cytosolic siRNA transfer on VAMP2-expressing cells without relying on lysosomal escape challenges the paradigm of conventional lipid nanoparticles. Future DFV designs could incorporate membrane components derived from viruses, immune/neuron/tumor cells, or extracellular vesicles to tailor fusogenicity and cellular specificity. Such hybrids may serve as precision vehicles for mRNA, protein, or gene-editing payloads, merging the structural fidelity of DNA nanotechnology with the functional sophistication of biological membranes. In this sense, DFV-mediated programmable fusion not only deepens our understanding of membrane mechanics but also offers a generalizable blueprint for engineering synthetic organelles and intelligent therapeutic nanotools.

## Acknowledgement

This work was financially supported by National Natural Science Foundation of China grants (22507076; 22277077; 32371283; 32571610; 22277017), the Key R&D Program of Shandong Province, China (2024CXPT029), the Taishan Scholars Program (2023000027), the Special Supporting Funds for Leading Talents at or Above the Provincial Level in Yantai City (2023000090), the Key Projects of Yantai Science and Technology Innovation Development Plan (2024JCYJ059), and Innovative research team of high-level local universities in Shanghai (SHSMU-ZLCX20212602). Fundamental Research Funds for the Central Universities (YG2025QNB37). We would like to thank for data collection in the Instrument Analysis Center (IAC), Shanghai Jiao Tong University on cryo-EM. We also show our great gratitude to all the special friends who supported our project and the authors.

## AUTHOR CONTRIBUTIONS

Q. Shi and Q. Yang performed most of the experiments, analyzed the data, arranged the figures, and prepared the manuscript. F. Li and Z. Wu provided all SNARE-related proteins, constructed the EGFP-VAMP expression plasmid, and characterized its cellular expression levels. M. Bao optimized the conditions and quantitative procedures for membrane-protein reconstitution. S.W. assisted with AFM experiments. K. Huang, and J. Liu assisted in structure assembly and cell culture. Y. Chen and Y. R. Yang contributed to cryo-EM data analysis and generated 2D-class images. Y. Wang and X. Bian participated in the initial conceptual discussion, structural design, and assembly strategy. Y. Yang and Z. Wu jointly conceived and supervised the project, interpreted the data, and finalized the manuscript.

## Supporting Information

Supplementary materials include experimental details; additional figures illustrating structural design, gel electrophoresis, fluorescence kinetics, nsTEM, cryo-EM, nFCM, confocal imaging, and flow cytometry results with corresponding quantitative or statistical analyses; as well as tables listing all DNA sequences, modifications, and lipid compositions used in this study. All supplementary materials are available online.

