## Supplementary material for "DNA Framework Nanoreactor for Programmable Membrane Fusion": Supporting Information.docx

1. **Methods**

**Materials and reagents**

The 7,560-nucleotide (nt) circular single-stranded DNA (ssDNA) scaffold (p7560, derived from M13mp18) was purchased from New England Biolabs (NEB). DNA staple and docking strands were synthesized and HPLC-purified by Sangon Biotech. Lipids were purchased from Avanti Polar Lipids. Recombinant VAMP2, t-SNARE complex (Syntaxin-1A/SNAP-25B), and the pEGFP-VAMP2 plasmid were provided by Wu Lab, as previously described (*1, 2*).

The flipped VAMP2/T27A construct was generated by fusing a bovine pre-prolactin signal peptide (MDSKGSSQKGSRLLLLLVVSNLLLCQGVVST), preceded by a Kozak sequence, to the 5’ end of the VAMP2/T27A coding sequence to invert its membrane topology. The T27A mutation was introduced to prevent glycosylation. The resulting fragment was cloned into pIRES2-EGFP or pcDNA3.4 vectors using standard molecular cloning, yielding plasmids designated flipped VAMP2/T27A–IRES2–EGFP and flipped VAMP2/T27A, respectively.

n-Octyl-β-D-glucopyranoside (OG), cytochalasin D, and other small-molecule inhibitors were obtained from Rhawn, MCE and Sigma-Aldrich. Silica nanosphere standards (S-16M-Exo) and fluorescent reference particles (QS2503) were obtained from NanoFCM.

**Assembly and purification of DNA soccer-ball framework (DSF)**

The truncated icosahedral DSF was designed using Tiamat software. Assembly was performed by mixing the p7560 scaffold (10 nM) with staple strands (100 nM) at a 1:10 molar ratio in 1× TE-Mg^2+^ buffer (5 mM Tris, 1 mM EDTA, 10 mM MgCl_2_, pH 8.0). Thermal annealing was carried out in a thermocycler (Applied Biosystems) with the following program: 95 °C for 5 min; 65 °C to 4 °C at -1 °C per 15 min (~15 h total). Assembled DSFs were purified by rate-zonal ultracentrifugation on a glycerol gradient. Structural integrity and morphology were verified by atomic force microscopy (AFM, Multimode 8, Bruker) in fluid mode using freshly cleaved mica as substrate. Purified DSFs were quantified using a Nanodrop One (Thermo Fisher) and stored at -20 °C.

**Preparation of proteoliposome mixtures**

Lipid stocks in chloroform were mixed in glass vials according to **Table S3** (total 1.25 µmol), dried under N_2_ for 30 min, and placed under vacuum overnight. The lipid film was resuspended in 500 µL of 1× hydration buffer (25 mM HEPES, 400 mM KCl, 10 mM MgCl_2_, pH 7.0) containing 1% (w/v) OG. For proteo-liposome preparation, VAMP2 (L/P = 200:1) or t-SNARE complex (L/P = 400:1) was added before resuspension. Samples were vortexed to mix thoroughly and stored at -80°C.

**Construction of DNA framework vesicles (DFVs)**

Cholesterol-modified DNA inner-anti handles (21 nt) were hybridized to the corresponding inner handles of the DSF (37 °C, 1 h) in hybridization buffer with 1% OG. To form DFVs, 100 µL of cholesterol modified DSF (20 nM) was mixed with 110 µL hydration buffer and 40 µL of lipid mixture (2.5 mM in OG), yielding a final OG concentration of 0.6%. The mixture was dialyzed overnight at 4 °C in 2 L hydration buffer using a Float-A-Lyzer device (8-10 kDa MWCO, Spectrum Labs). The sample was subjected to iodixanol density gradient ultracentrifugation (0–24%, 50,000 rpm, 5 h). Fractions (200 µL each) were collected and analyzed by SDS-agarose gel electrophoresis (SDS-AGE); DFV-containing fractions were pooled and stored at 4 °C (**Figure S2**).

**Quantification of SNARE protein incorporation**

DFV samples (5 nM, 30 µL) and protein standards (250 ng VAMP2 or t-SNARE) were mixed with 5× SDS loading buffer, heated at 100 °C for 15 min, and separated on 15% SDS–PAGE gels. Proteins were visualized using a silver staining kit (Beyotime) and band intensities were quantified in ImageJ to estimate the average protein copy number per DFV.

**SDS–agarose gel electrophoresis (SDS–AGE)**

DFV samples were electrophoresed in 1.5% agarose gels containing 0.05% SDS. Gels were run at 65 V for 100 min in running buffer (0.5× TBE, 10 mM MgCl_2_, 0.1% SDS) in an ice bath. Gels were visualized and documented using a ChemiDoc™ imaging system (BIO-RAD).

**DFV nanoreactor assembly**

All DFV variant monomers were quantified by SDS-AGE using a DSF standard as reference (10 nM). DFV-A and DFV-B carrying specific linkers were mixed with stoichiometric ratios to assemble dimers (H/C, H/F or P/C; 1:1), trimers (H/C or P/C; 1:2), pentamers (H&P/C; 1:2:2) and linear arrays (H/C or P/C; 1:1). All assemblies were incubated at 37 °C for 6 h (for kinetics) or 18 h (for nsTEM and cryo-EM imaging). Assembly yields were quantified from SDS–AGE band intensities using ImageJ.

**Nano-flow cytometer (nFCM)**

DFV-A*_VAMP2_* and DFV-B*_tSNAREs_* were fluorescently labeled by incorporating FITC- or Cy5-conjugated cholesterol, respectively, during liposome formation (**Table S3**). Single-particle analysis was performed using a Flow NanoAnalyzer (nFCM, U30E) equipped with 488 nm (FITC: 525/40 nm filter) and 638 nm (Cy5: 670/30 nm filter) lasers. Silica nanosphere standards (S-16M-Exo, NanoFCM) and fluorescent reference particles (QS2503, NanoFCM) were used for size calibration and concentration quantification. Samples were analyzed at an operating pressure of 1.0 kPa following a 60 s pre-run stabilization. Data were acquired for 60 s per sample, with signal thresholds set automatically. Sample concentrations were determined by subtracting counts from 1× hydration buffer blanks. All data were processed using NF Profession 2.0 software and FlowJo_v10.8.1 software.

**Electron microscopy**

For negatively stained TEM imaging, 5 µL of sample (5 nM) was applied to glow-discharged carbon-coated grids, stained with 2% uranyl acetate, and imaged on a Hitachi HT7700. For cryo-EM, 4 µL of sample (15 nM) was applied to Lacey carbon grids, blotted for 3 s at 22 °C and 100% humidity, and plunge-frozen in liquid ethane using a Vitrobot Mark IV (Thermo Fisher Scientific). Imaging was performed on a Talos L120C G2 (Thermo Fisher Scientific) and further analyzed using Image J software.

For the analysis of fusion intermediates (**Figure S8**), particles were manually picked from cryo-EM micrographs and subjected to reference-free 2D classification in CryoSPARC (*3*). Representative 2D class averages were rotationally aligned to match schematic diagrams using a custom Python script. All images were preprocessed using ImageJ software.

**Fusion stage quantification**

Cryo-EM images were analyzed using ImageJ. Distances between the inner leaflets of the two vesicles (D_in_) were measured for Stages 1-3; while outer diameters of the lateral dilated pore after fusion (D_out_) were measured for Stages 4-6. Histograms were generated in Origin and fitted to multi-peaks Gaussian distributions.

**Lipid mixing and rounds of fusion (ROF) assays**

Membrane fusion was monitored by mixing R18- or NBD/Rhodamine-labeled DFVs with unlabeled partner DFVs at specified stoichiometries in black 96-well plates (final volume: 100 µL). Fluorescence was recorded every 2 min for 6 h at 37 °C (λ_ex_/λ_em_ = 560/590 nm for R18; 460/535 nm for NBD/Rhodamine) using a Synergy H1 microplate reader (BioTek). After 6 h, 25 µL of 20% Triton X-100 was added to determine maximum fluorescence.

ROF values were calculated from calibration curve of labeled-to-unlabeled liposome mixtures (ratios 1:0 to 1:8). All fluorescence signals were background-corrected by subtracting the DFV-A only or 1:0 values prior to detergent addition and normalized to the detergent-induced maximum.

*Technical Note on R18 dequenching assay*
Although R18-based lipid mixing assays can sometimes be affected by dye transfer between membranes or nonlinear fluorescence responses, these effects were minimized here by using a low R18 labeling density (5 mol%), including protein-free and detergent-solubilized controls, and establishing a calibration curve from defined mixtures of labeled and unlabeled lipids (ratios 1:0 to 1:8). This allowed fluorescence recovery to be quantitatively converted into rounds of fusion (ROF), following the framework established by Parlati *et al (4)*. Under these controlled conditions, the R18 dequenching assay provided a robust and reproducible readout of membrane merger.

**Cell culture and transfection**

HeLa cells were cultured in DMEM supplemented with 10% FBS and 1% penicillin–streptomycin at 37 °C in a humidified incubator with 5% CO₂. For surface VAMP2 expression, cells were transfected with pEGFP-VAMP2 (flipped orientation) using Lipofectamine 2000 (Thermo Fisher Scientific) following the manufacturer’s instructions. Assays were performed 24 h post-transfection.

**Cell-surface protein biotinylation**

To confirm plasma membrane localization of flipped VAMP2, a surface biotinylation assay was performed 24 h post-transfection. Transfected HeLa cells were placed on ice and washed with ice-cold PBS^++^ (PBS supplemented with 1 mM CaCl_2_ and 0.5 mM MgCl_2_, pH 7.4). Cells were then incubated with 1 mL of Sulfo-NHS-SS-Biotin (1 mg/mL) at 4 °C for 30 min with gentle agitation. Unreacted biotin was quenched by washing with 100 mM glycine (one quick rinse followed by two 5-min washes, all at 4 °C). Cells were subsequently washed three times with PBS^++^, lysed in 700 µL RIPA buffer containing protease inhibitors, and incubated for 20 min at 4 °C with gentle shaking. Lysates were sonicated and clarified by centrifugation, and the resulting supernatants were incubated with 40 µL avidin–agarose beads (1:1 slurry) for 2 h at 4 °C with rotation. The beads were collected by centrifugation, and the bound biotinylated proteins were eluted and analyzed by Western blotting.

**Endocytosis inhibition and live-cell fusion assay**

Cells were pretreated with 2 µM cytochalasin D (or other inhibitors as shown in **Figure S27**) for 30 min. DFV-A (0.5 nM, 5% R18) was then added and incubated for 3 h at 37 °C. Cells were washed, fixed, and imaged by confocal microscopy. Mean fluorescence intensity (MFI) of R18 at the plasma membrane was quantified using ImageJ.

**siRNA encapsulation and delivery**

siEGFP was hybridized to cholesterol-anchored DNA strands through a three-strand bridge (**Table S4**) and co-reconstituted into DFV-A*_t-SNARE_* during vesicle formation. Loaded DFVs were purified using iodixanol density gradient ultracentrifugation. For +RNase A samples, DFVs were treated with 3U/µL RNase A for 30 min at 37 °C to digest externally associated siRNA. Following digestion, DFVs were purified through Amicon Ultra-0.5 mL centrifugal filters (100 kDa, Millipore) to remove RNase A and cleaved siRNA fragments, then resuspended to match the DFV concentration of untreated controls (-RNase A). EGFP knockdown efficiency was quantified by flow cytometry 48 h post-incubation under both cytochalasin D-treated (plasma membrane-fusion specific) and standard culture conditions.

**Statistical analysis**

Statistical analyses were performed using GraphPad Prism 9.0. All data in this study are presented as mean ± SD. Statistical significance was determined using *P* values calculated by two-way analysis of variance (ANOVA) and paired Student’s t-test. Significance levels are indicated as follows: NS (not significant), **P* < 0.05, ***P* < 0.01, and ****P* < 0.001.

2. **Supplementary Figures**

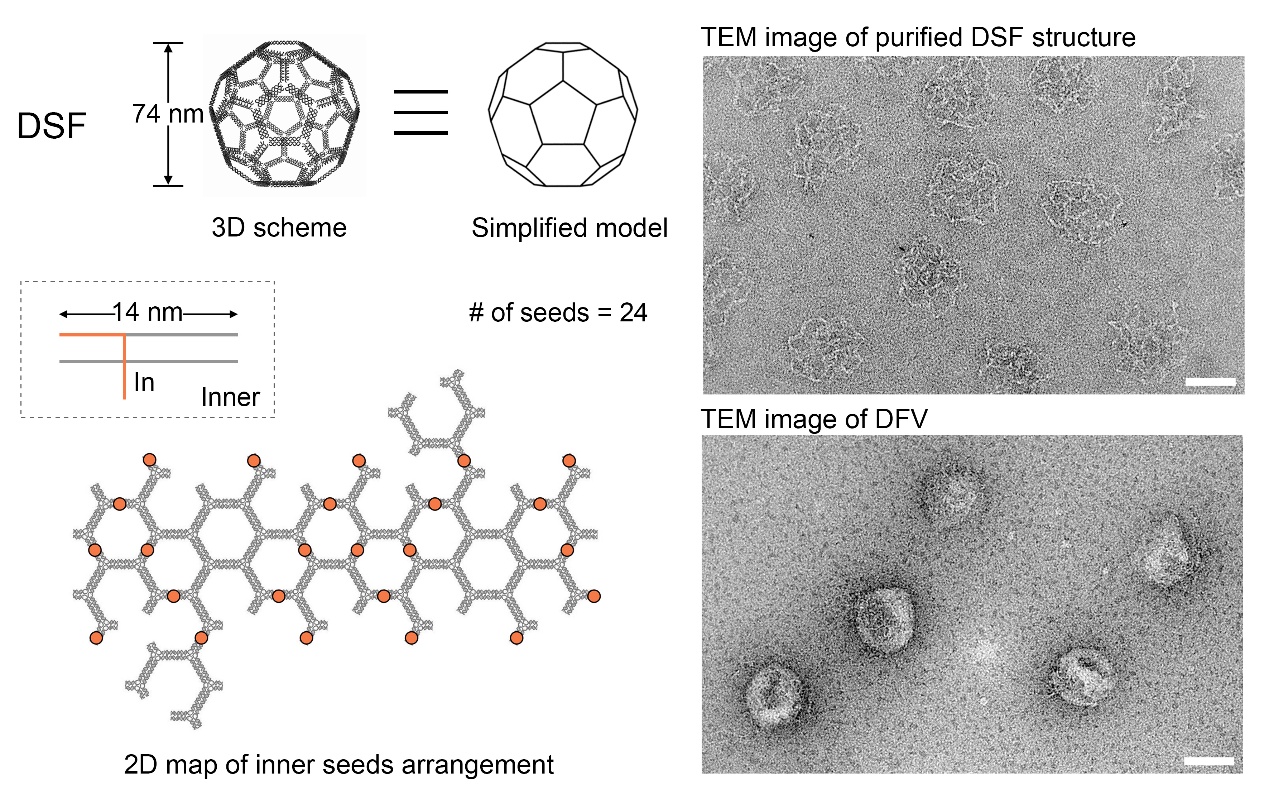

**Figure S1. Schematic diagrams of DSF and positions of inner seeds.**
3D schematic representation of DSF alongside a simplified model, with cholesterol modification sites indicated by orange dots on the 2D DSF blueprint (left panel). Representative TEM image of purified DSFs and DFVs show the highly uniform and intact of the structures (right panel). Scale bar: 50 nm.

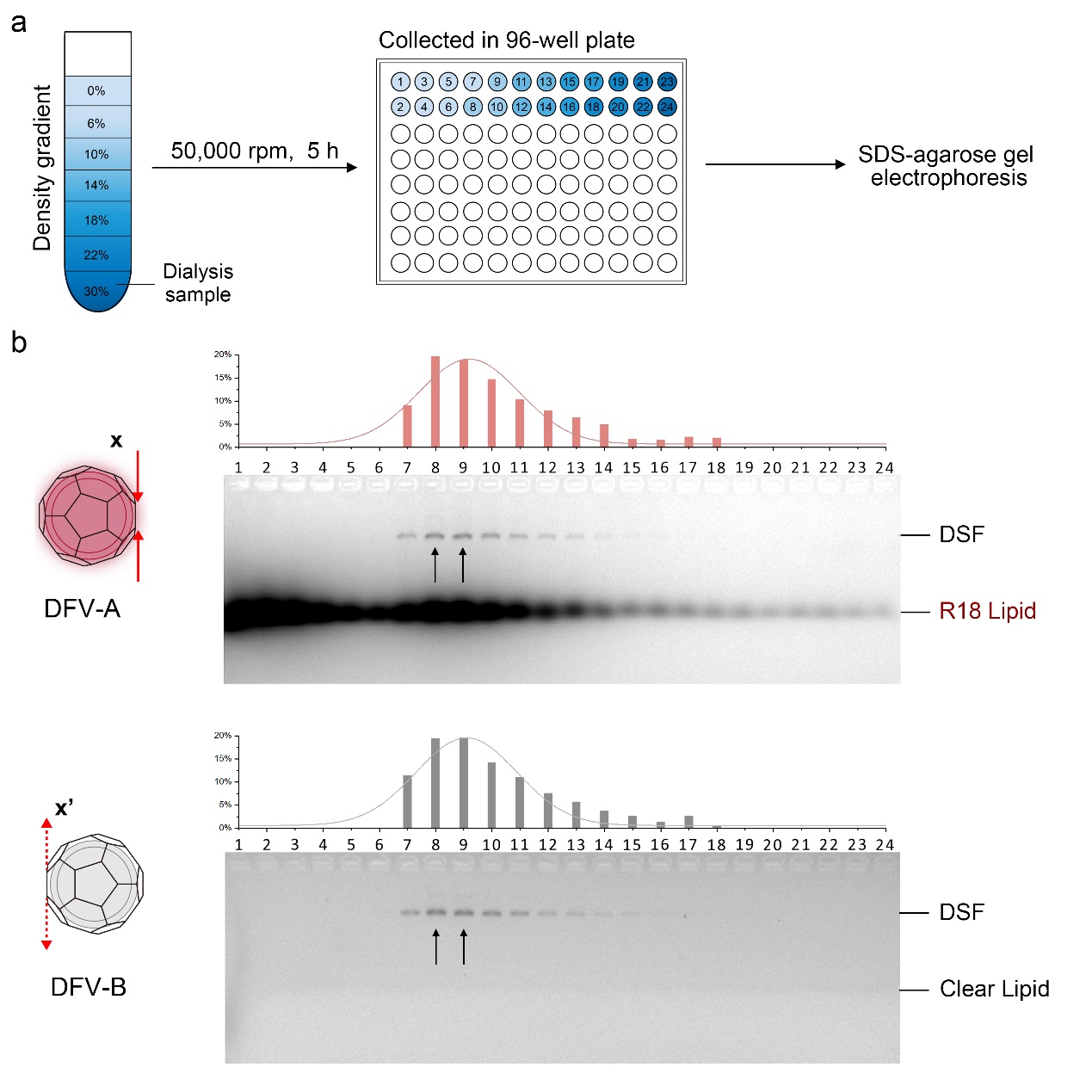

**Figure S2. Purification and collection of DFV-A and DFV-B.**
(a) Workflow of DFV purification from dialysis samples by iodixanol density gradient centrifugation (0-24%, w/v). (b) SDS-agarose gel electrophoresis showing product distribution, with peak fractions (black arrows) collected for DFV-A (F8 + F9) and DFV-B (F8 + F9), respectively. The DSF structure was labeled with TAMRA fluorescence.

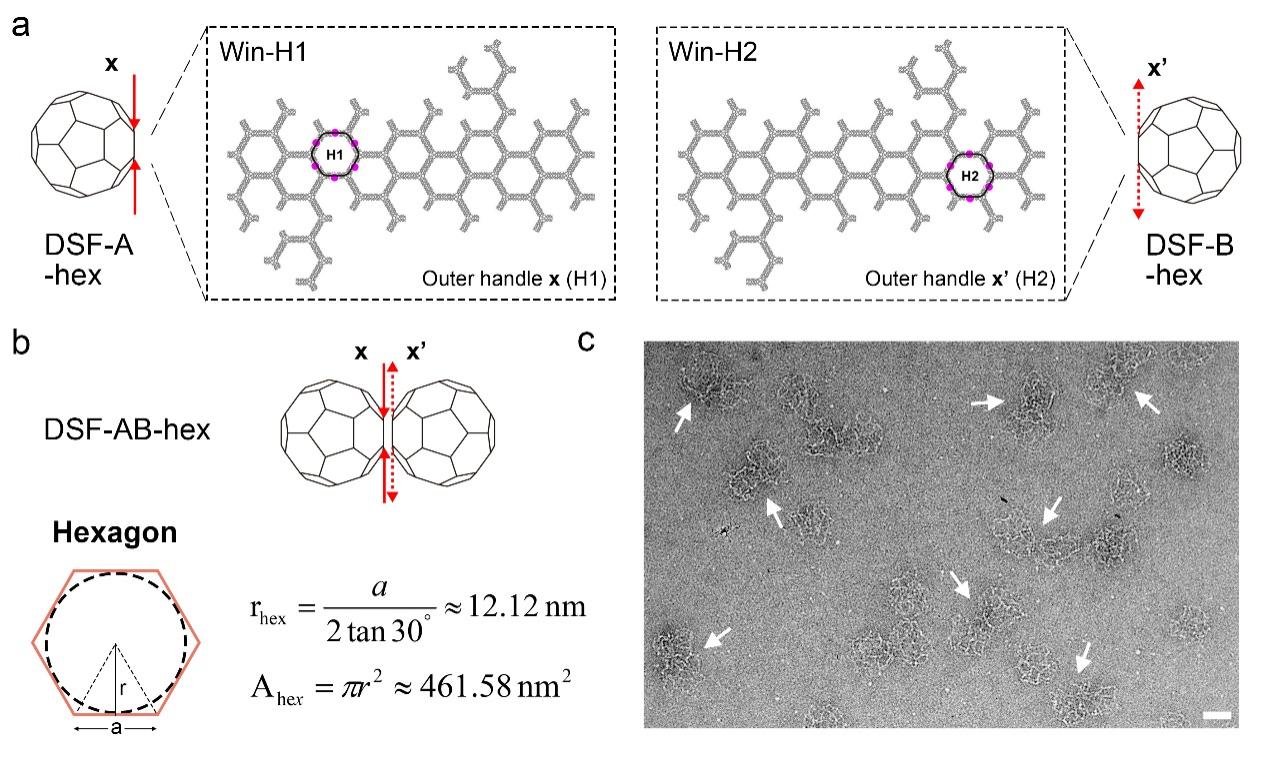

**Figure S3. Design and characterization of the hexagonal docking window.**
(a) Schematics of the selected hexagonal windows for tethering: Win-H1 on DSF-A (left) and Win-H2 on DSF-B (right). Purple dots indicate the extension sites of the complementary DNA linker strands. (b) Calculated geometric parameters of the hexagonal window including the radius ($r_{\text{hex}}$) and area ($A_{\text{hex}}$) of the inscribed circle. (c) Representative TEM image showing the successful dimerization of DSF-AB (white arrows) via hexagonal window docking. Scale bar: 50 nm.

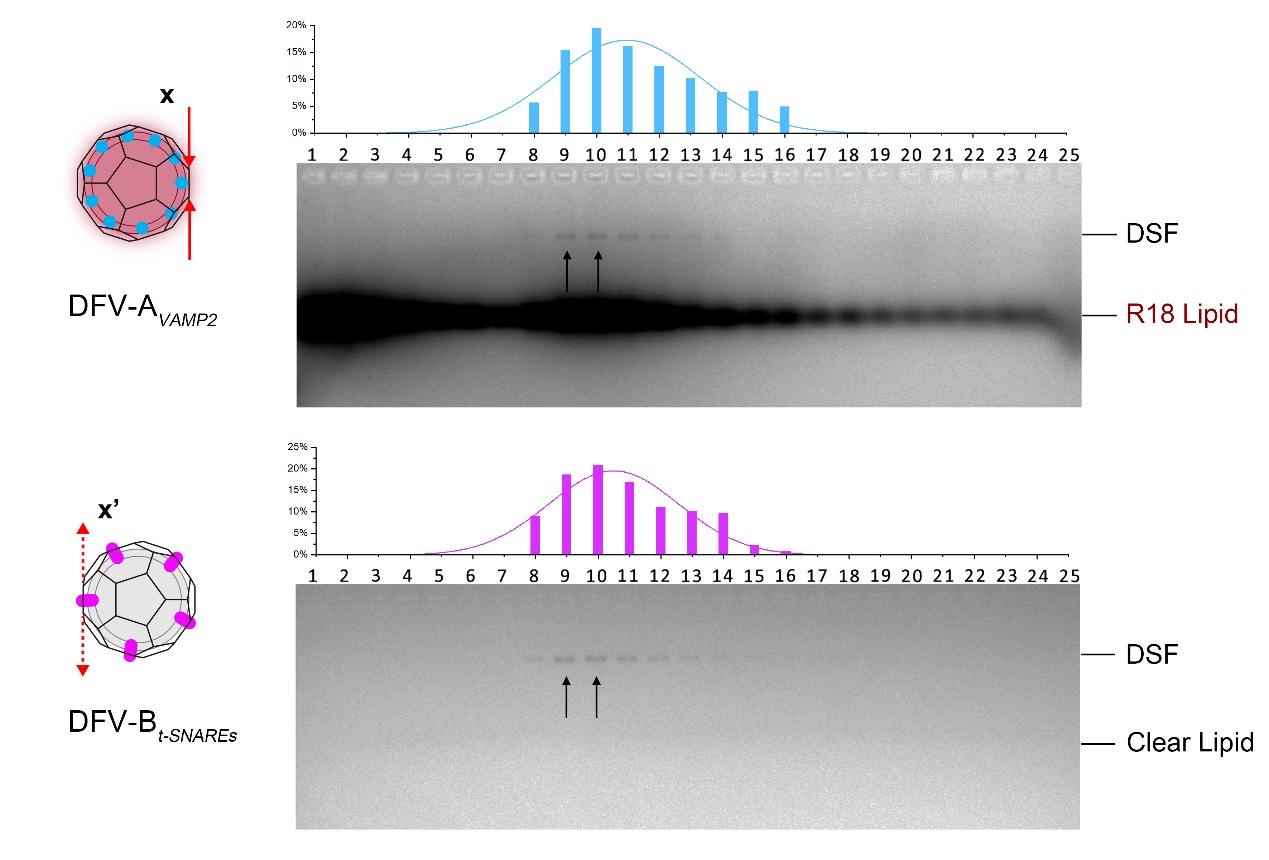

**Figure S4. Purification and collection of proteo-DFVs.**
SDS-AGE images illustrate the distribution of products following density gradient purification. The peak fractions (black arrows) for DFV-A*_VAMP2_* (F9 + F10) and DFV-B*_t-SNAREs_* (F9 + F10) were collected, respectively.

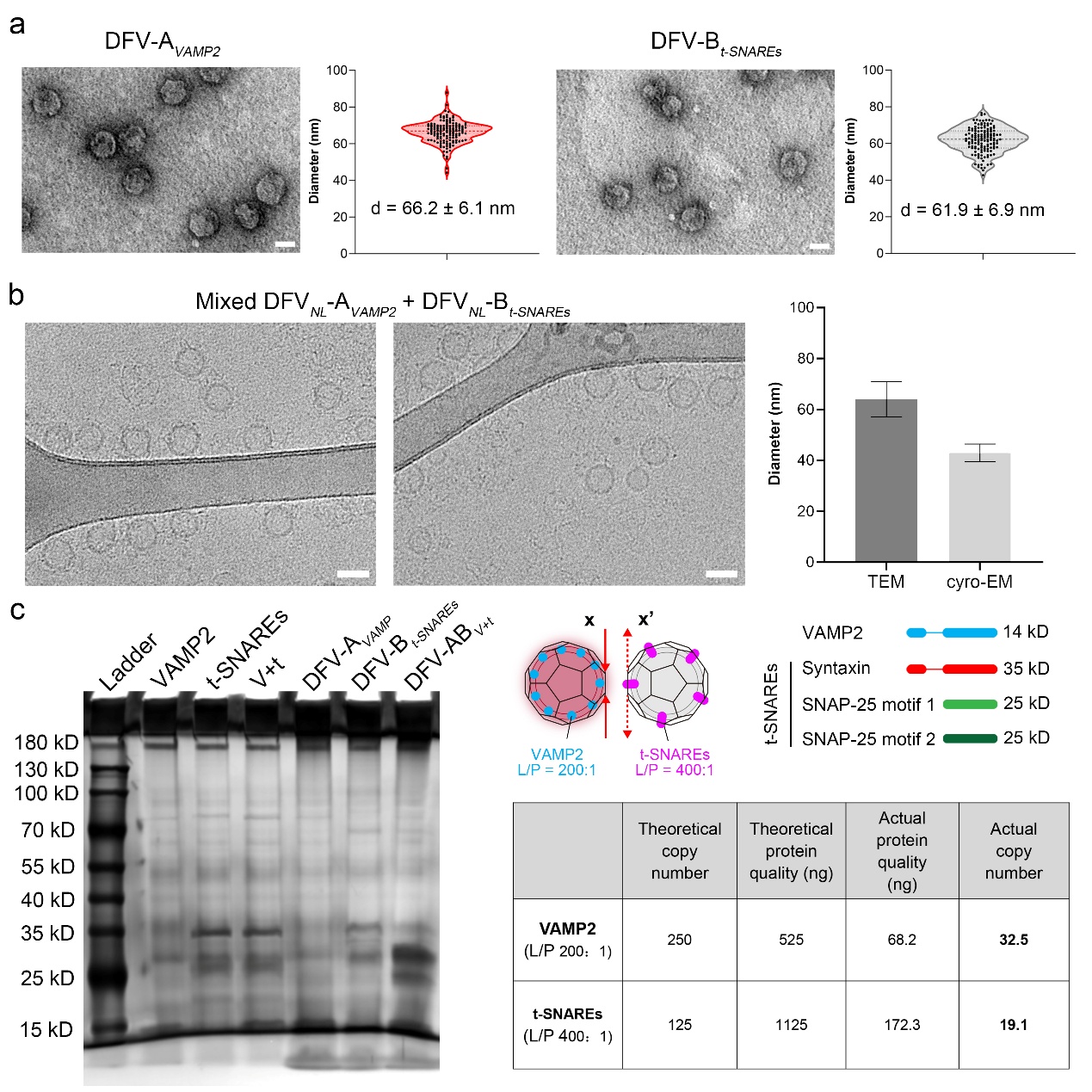

**Figure S5. Size distribution and protein quantification of proteo-DFVs.** 
(a) Representative nsTEM images of DFV-A*_VAMP2_* and DFV-B*_t-SNAREs_* with corresponding diameter distributions (mean ± SD). Scale bar, 50 nm. (b) Cryo-EM images of mixed DFV-A*_VAMP2_* and DFV-B*_t-SNAREs_* (note: without linkers), showing monodisperse vesicles with a diameter of 43.0 ± 3.5 nm, *n* = 205, compared with its nsTEM measurements (64.0 ± 6.9 nm, *n* = 280) in histogram panel. Scale bar, 50 nm. (c) Silver-stained SDS-PAGE of VAMP2, t-SNARE, and preassembled VAMP2–t-SNARE complex (V+t), alongside proteo-DFV samples. A schematic depicts subunit molecular weights, and the accompanying table summarizes protein copy numbers per DFV derived from band intensity calibration against known standards.

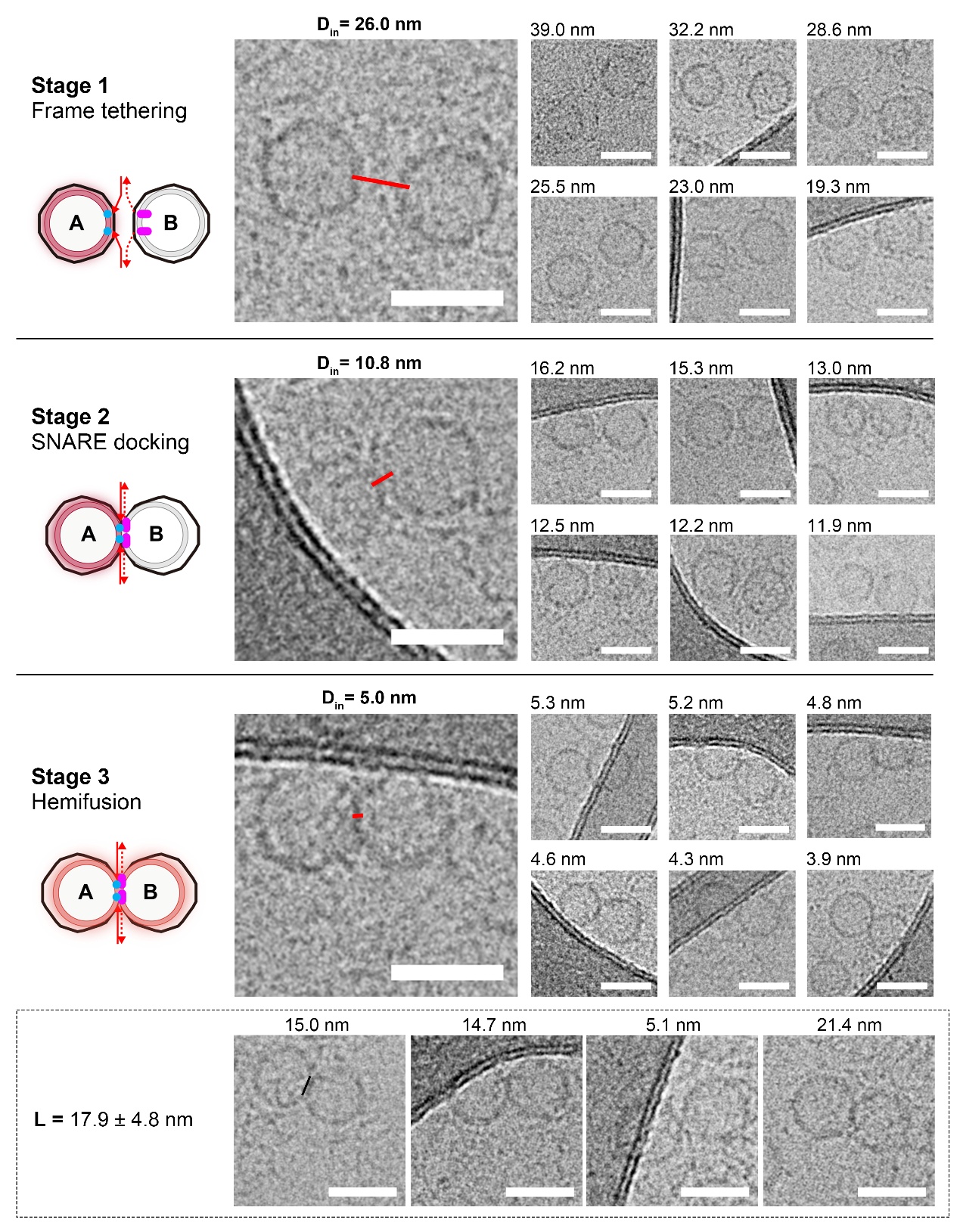

**Figure S6. Cryo-EM images of intermediates from Stage 1 to Stage 3 in DFV-AB.**
Representative cryo-EM images and corresponding measurements of **D_in_** for (**Stage 1**) frame-tethering, (**Stage 2**) SNARE docking and (**Stage 3**) hemifusion. Red lines indicate measurement positions. The length (**L**) of the hemifusion diaphragm in Stage 3 is also shown (mean ± SD; *n* = 39). Scale bar: 50 nm.

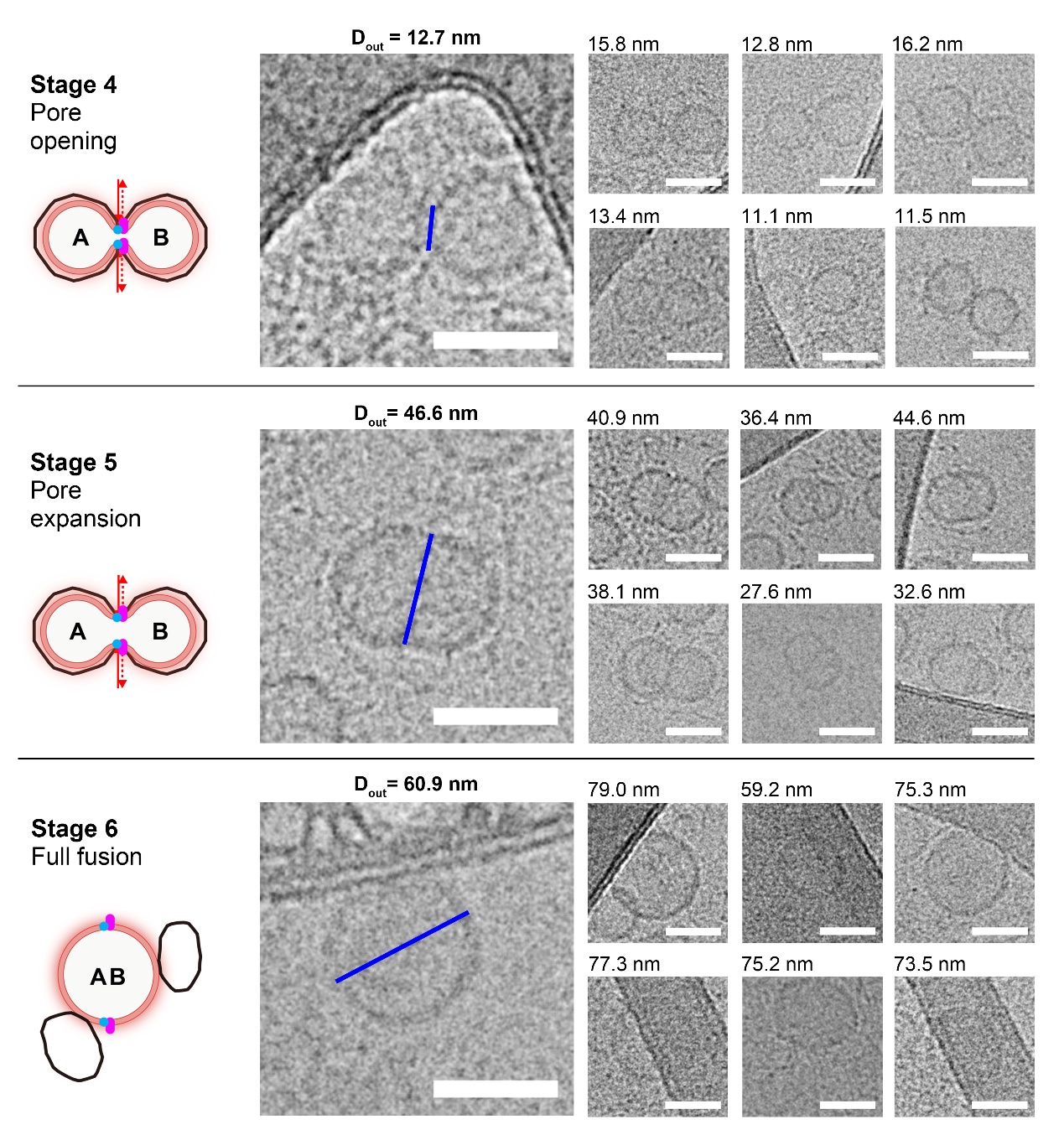

**Figure S7. Cryo-EM images of intermediates from Stage 4 to Stage 5 in DFV-AB.**

Representative cryo-EM images and corresponding measurements of **D_out_** for (**Stage 4**) pore opening, (**Stage 5**) pore dilation, and (**Stage 6**) Full fusion. Blue lines indicate measurement positions. Scale bar: 50 nm.

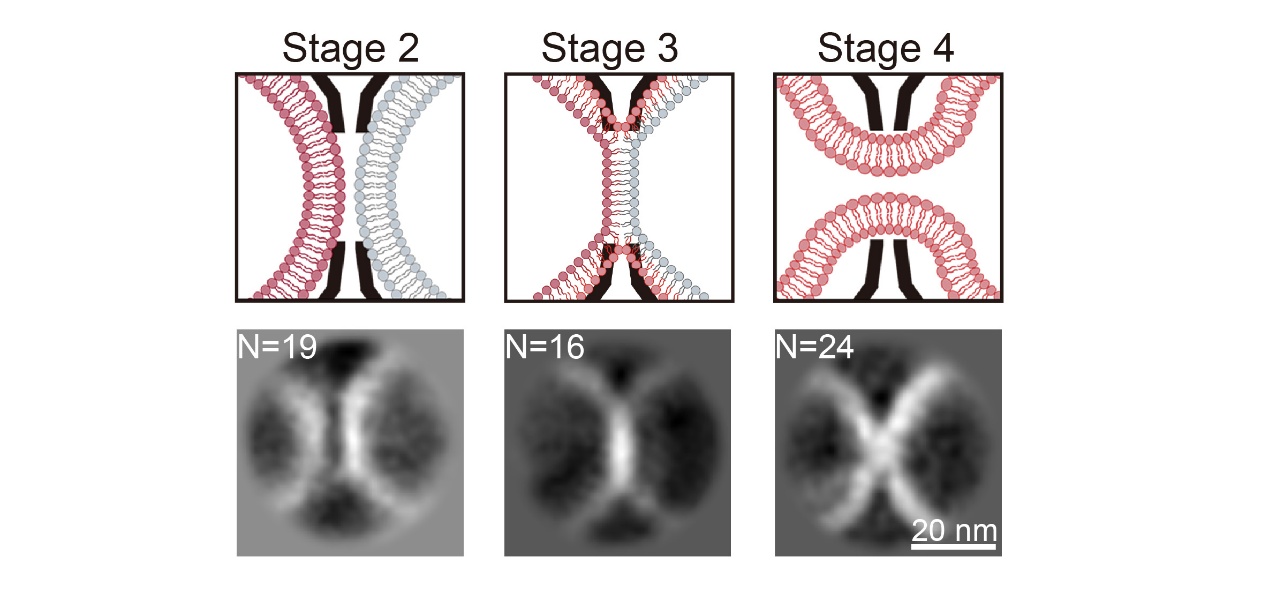

**Figure S8. Representative 2D class averages of SNARE-mediated membrane fusion intermediates.**
Schematics and representative 2D class averages for Stage 2, 3, 4 with selected particle numbers (N) shown. Scale bar, 20 nm.

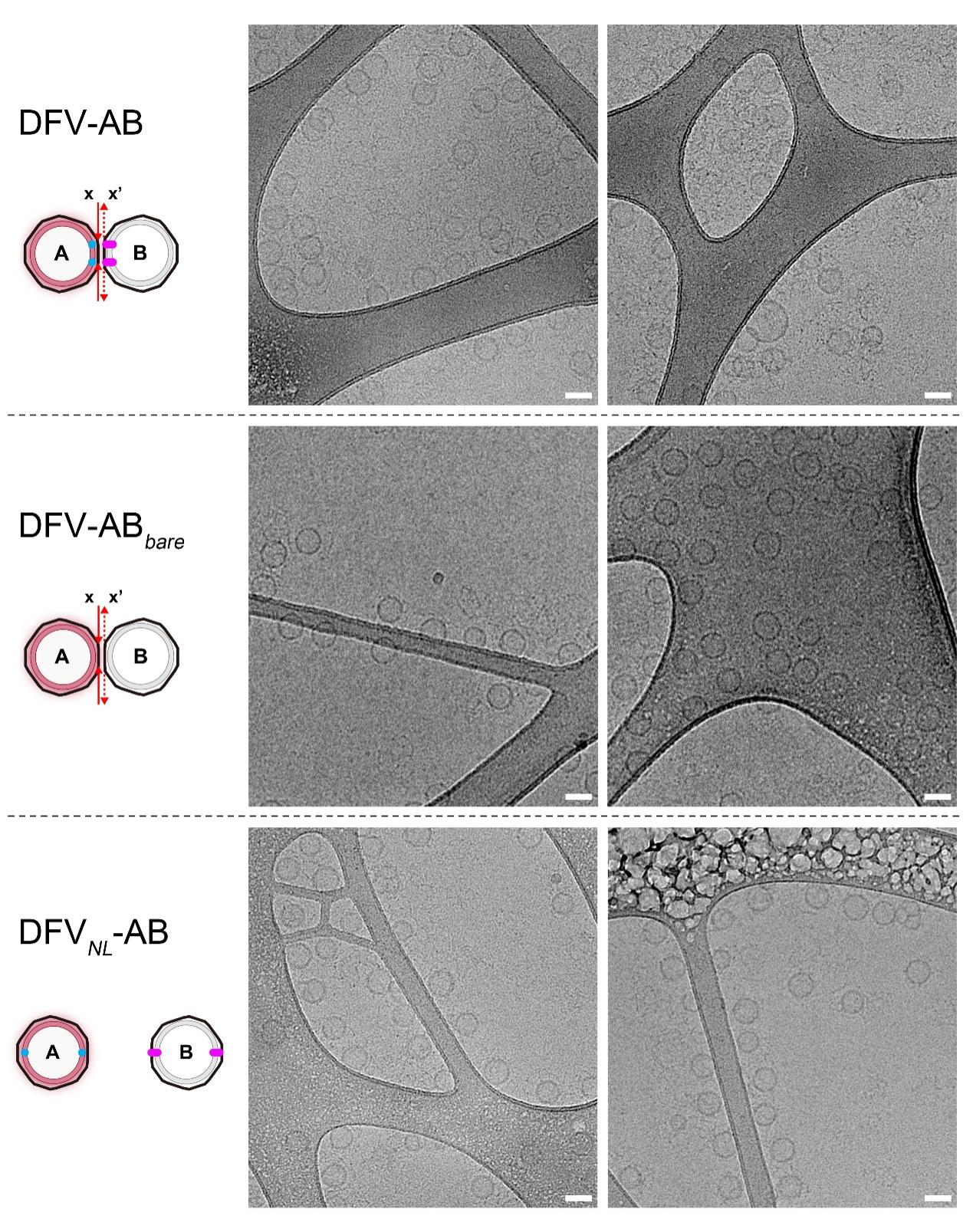

**Figure S9. Cryo-EM comparison of DFV-AB dimers under different tethering conditions.**
Representative cryo-EM images of *(top)* DFV-AB with SNARE proteins and DNA linkers, *(middle)* DFV-AB*_bare_* with DNA linkers but lacking SNARE proteins, and *(bottom)* DFV*_NL_*-AB lacking inter-vesicle linkers. Scale bar: 50 nm.

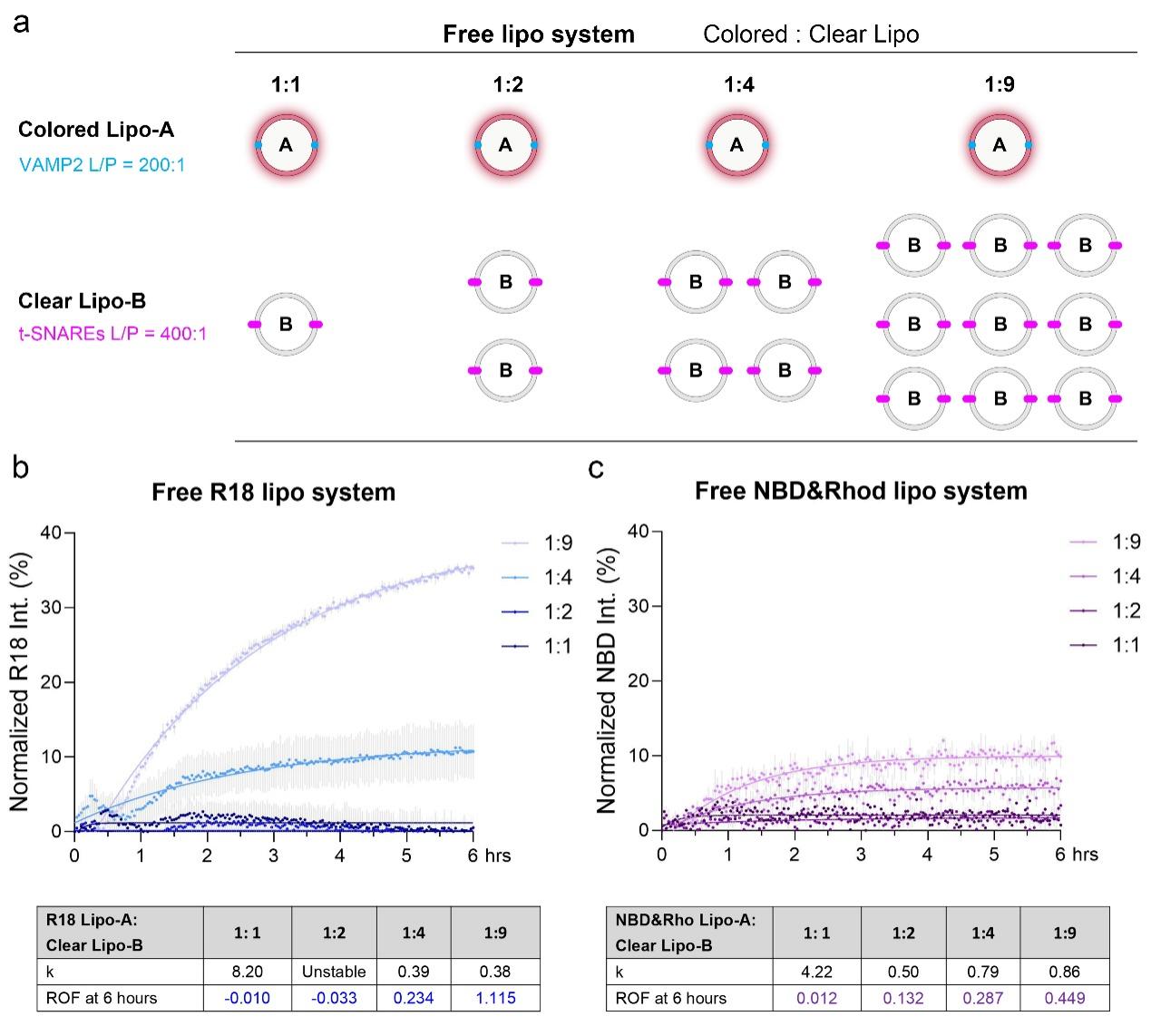

**Figure S10. Free liposome membrane fusion system.**
(a) Schematic illustration of the free liposome membrane fusion assay, showing the mixing ratios of colored Lipo-A to clear Lipo-B at varying ratios (1:1 to 1:9). (b) Normalized R18 dequenching kinetics of 5% R18 Lipo-A*_VAMP2_* fused with clear Lipo-B*_t-SNAREs_* at the indicated ratios. The accompanying table summarizes rate constants (k) and rounds of fusion (ROF) after 6 h. Corresponding NBD/Rhodamine fluorescent dequenching kinetics reflecting reduced FRET efficiency using 1.5% dual-labeled Lipo-A*_VAMP2_* and clear Lipo-B*_t-SNAREs_* are shown in parallel, with kinetic parameters tabulated below.

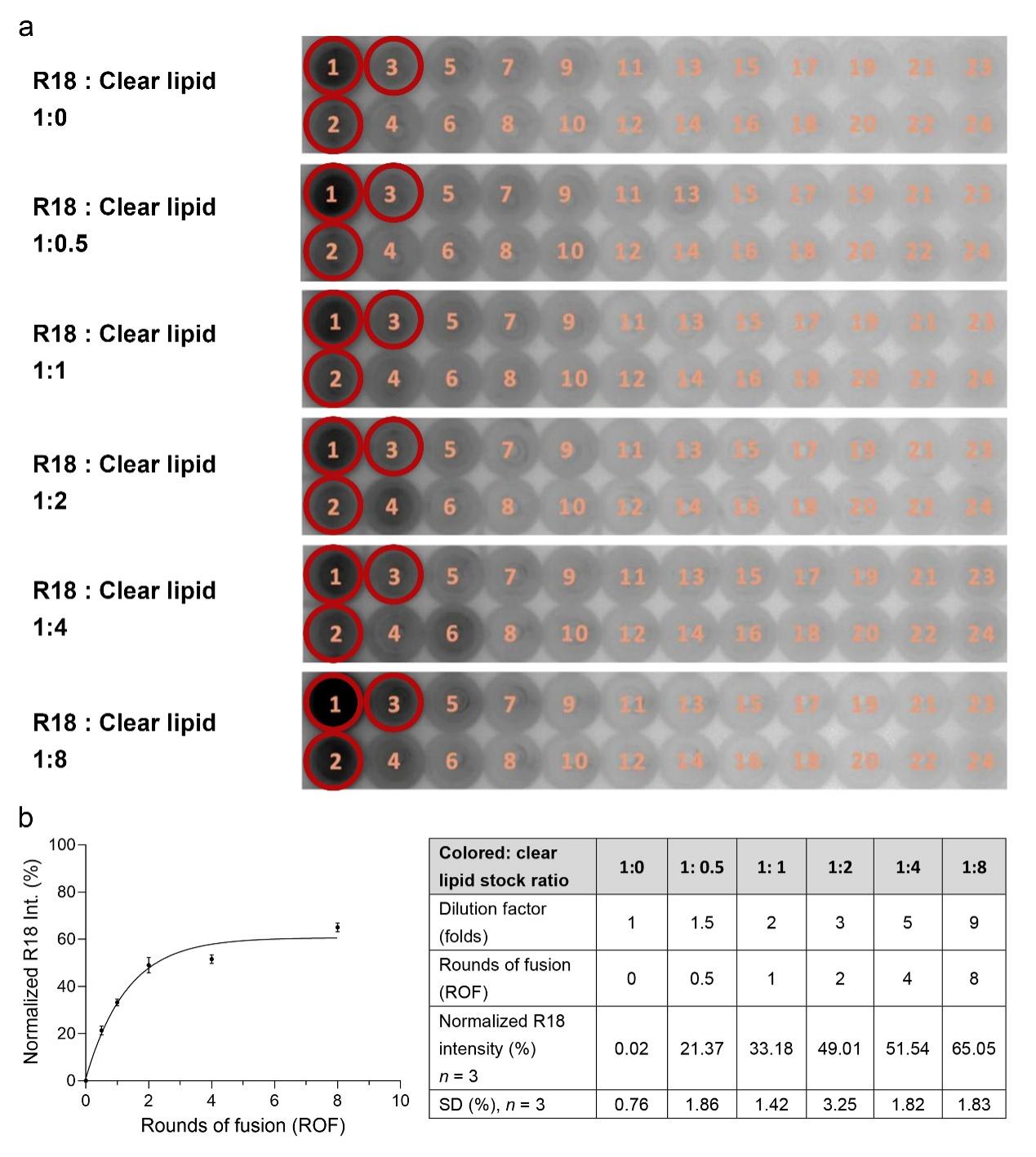

**Figure S11. Calibration curve for calculating ROF in the R18 system.**
(a) Mixtures of 5% R18 lipid and clear lipid were prepared from lipid stock at defined ratios: 1:0, 1:0.5, 1:1, 1:2, 1:4, and 1:8. Peak fractions (F1-F3), indicated by red circles, were combined to construct the calibration curve. (b) Calibration curve showing the relationship between R18 intensity and calculated rounds of fusion (ROF). Data points (mean ± SD, *n* = 3; as summarized in the right table) are fitted with a one-phase association model (black curve) for ROF determination.

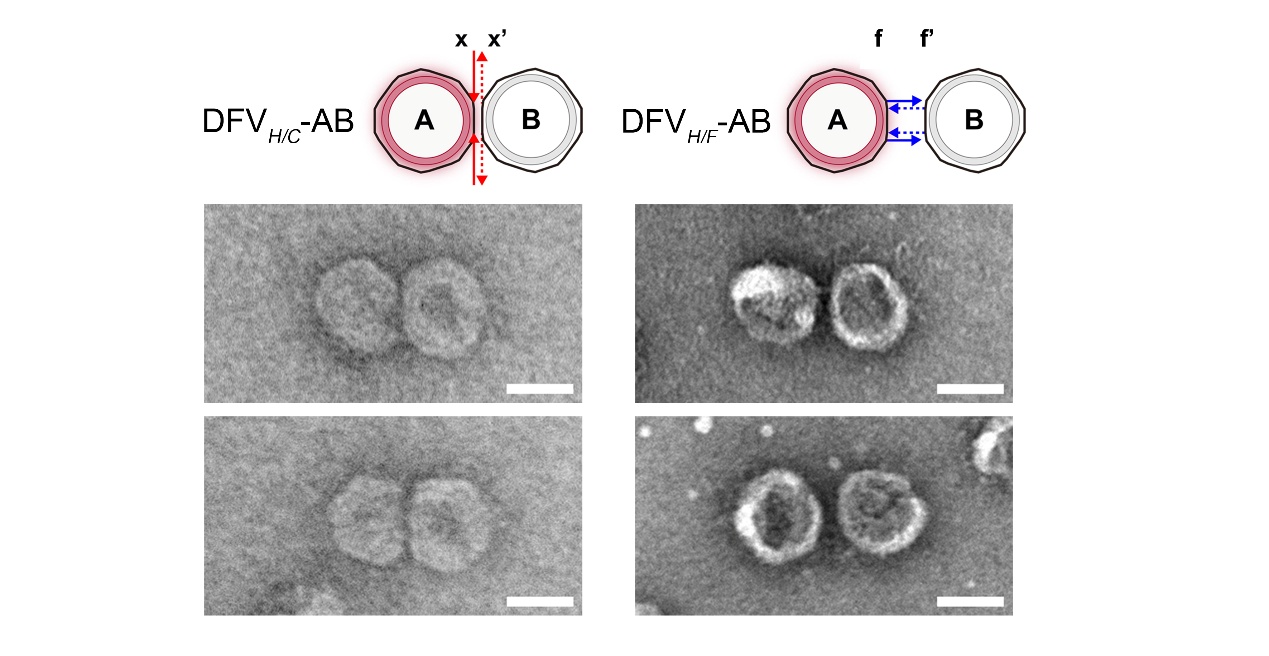

**Figure S12.** **Structural details and characterization of DFV-AB close and far constructs without SNARE proteins.**
Schematics and representative nsTEM images of DFV*_H/C_*-AB (linker: x/x’) and DFV*_H/F_*-AB (linker: f/f’) without protein reconstitution. Scale bar, 50 nm.

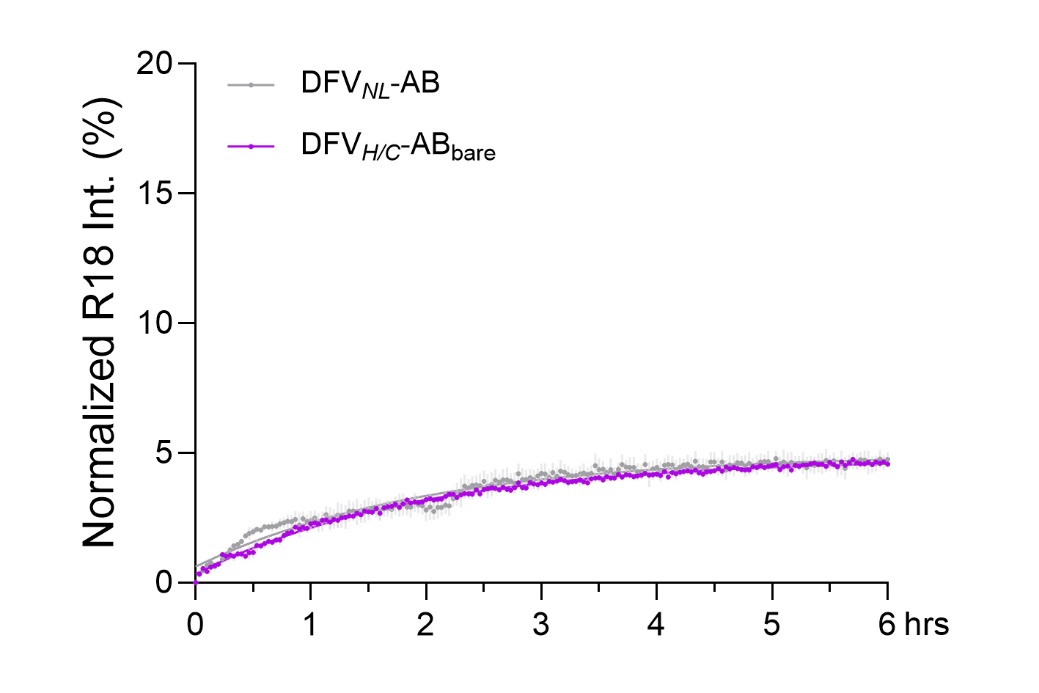

**Figure S13. Fusion kinetics of DFV-A+ DFV-B controls lacking SNARE proteins or DNA linkers.**
Time-dependent R18 dequenching curves of DFV*_H/C_*-AB*_bare_* (lacking SNAREs, purple) and DFV*_NL_*-AB (lacking inter-frame linkers, gray). The DFV*_NL_*-AB trace is consistent with the data presented in Figure 3a. The overlapping traces with minimal fluorescence recovery confirm the essential role of both SNARE proteins and programmed tethering of DNA frames in driving efficient membrane fusion.

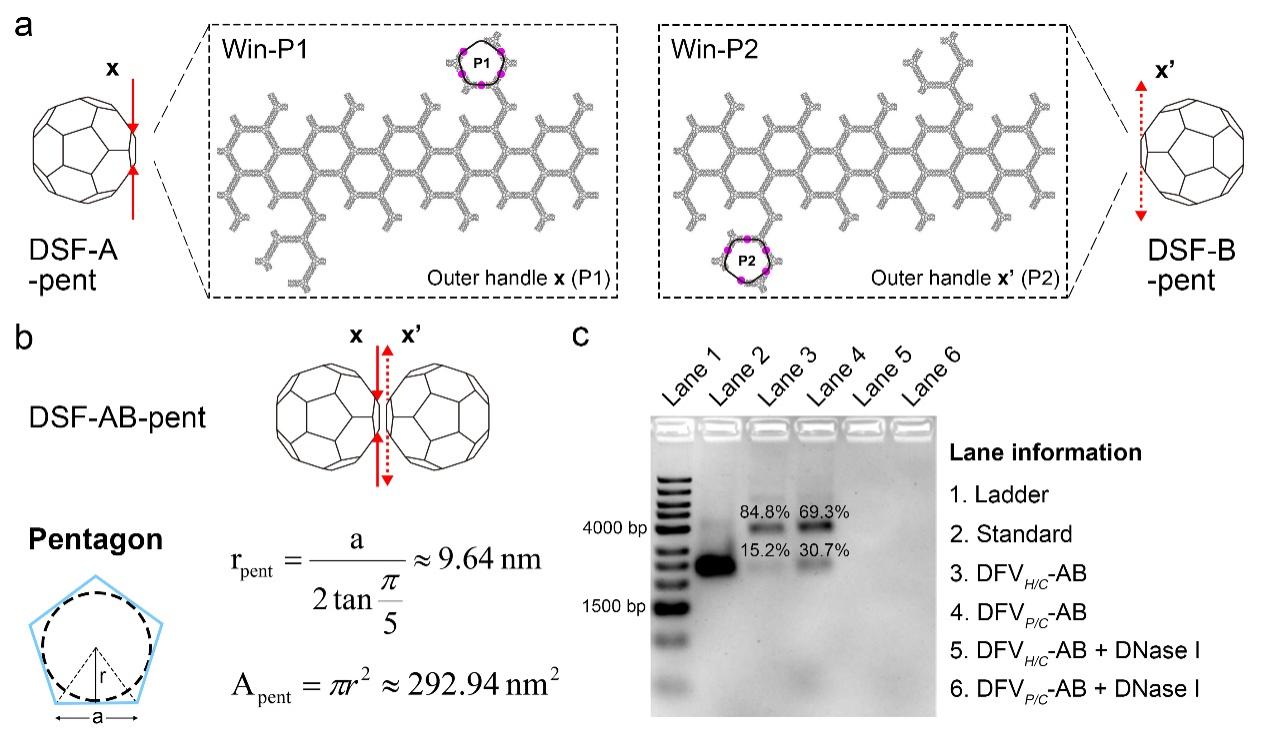

**Figure S14. Design and characterization of the pentagonal docking windows.**
(a) Schematics of pentagonal faces selected for DSF-A-pent (Win-P1) and DSF-B-pent (Win-P2). Purple dots indicate the linker extension sites. (b) Calculated geometric parameters of the pentagonal window including the radius ($r_{\text{pent}}$) and area ($A_{\text{pent}}$) of the inscribed circle. (c) SDS-AGE analysis of dimerization efficiency, before and after DNase I digestion. Percentage of corresponding bands were labeled.

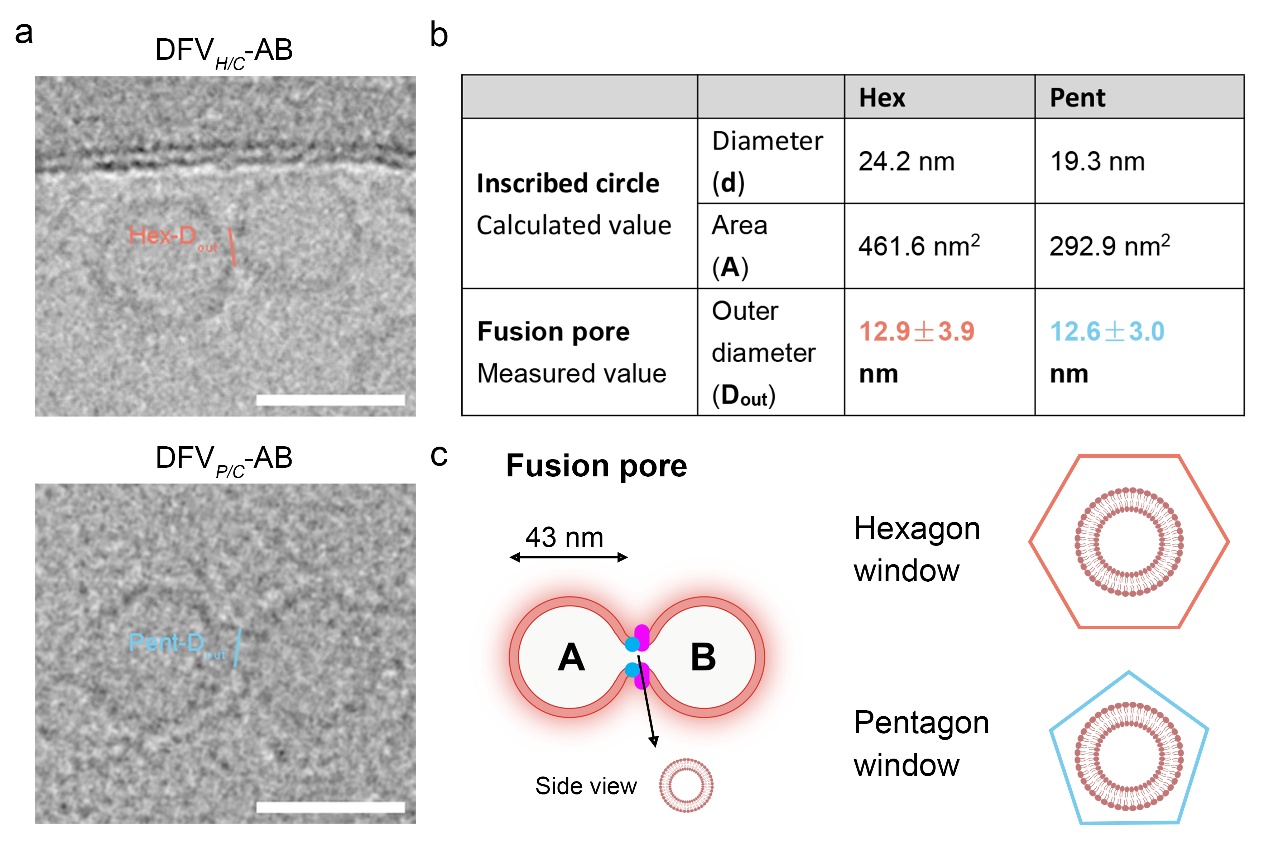

**Figure S15. Fusion pores in hexagonal vs. pentagonal docking windows.**
(a) Representative cryo-EM images and corresponding schematic cross-sections of fusion pores in DFV*_H/C_*-AB (hexagonal, *top*) and DFV*_P/C_*-AB (pentagonal, *bottom*). Scale bar: 50 nm. (b) Comparison of geometric parameters of hex. and pent. windows and measured fusion pore outer-diameters (**D_out_**, mean ± SD), showing conserved pore size despite differing window areas. (c) Structural model of fusion pore formation within DNA framework-confined membranes.

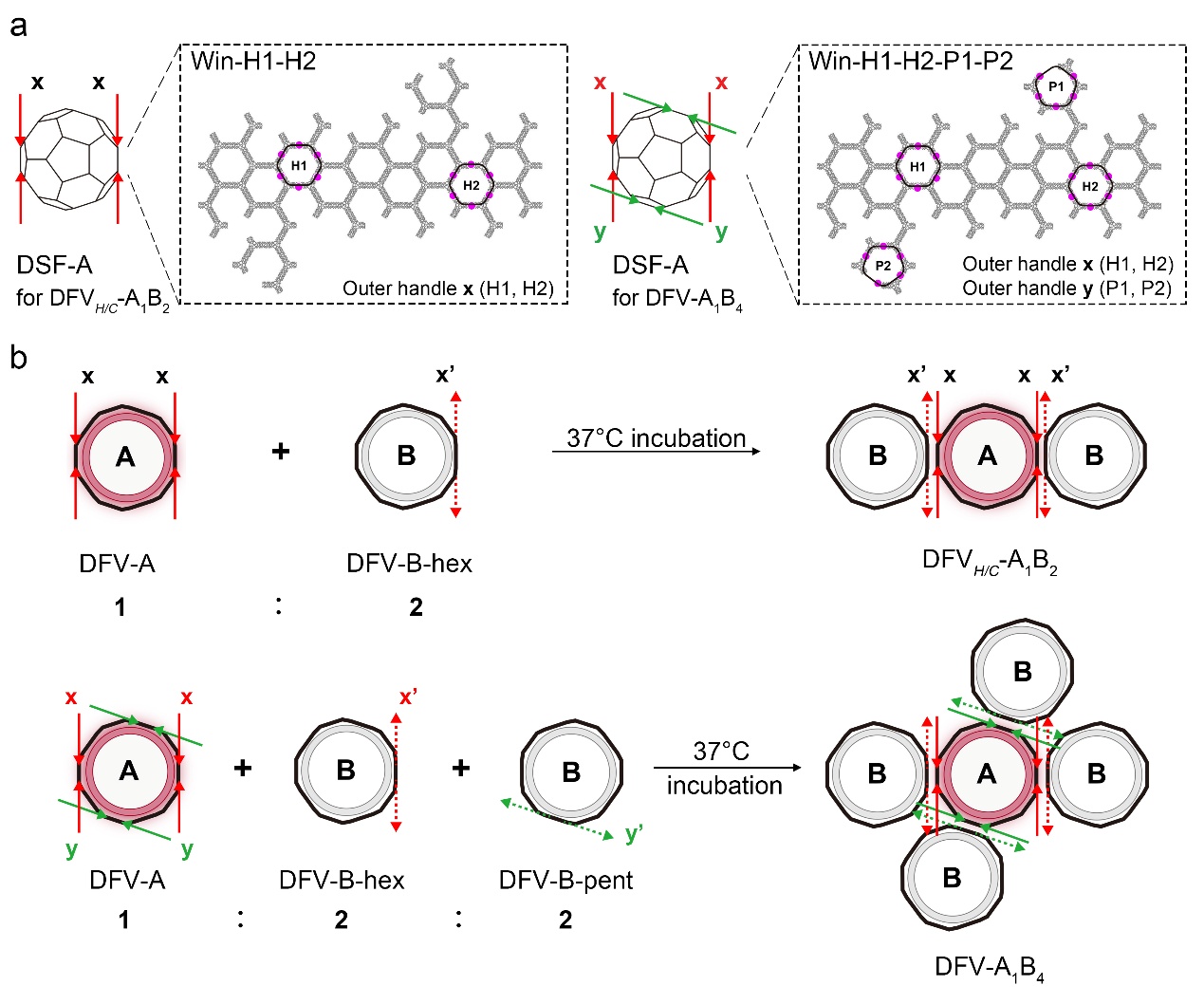

**Figure S16. Design and assembly strategy for DFV-A_1_B_2_ and DFV-A_1_B_4_ complexes.**

(a) Schematic design and corresponding 2D maps showing the docking windows and linker pairs used for DFV-A_1_B_2_ and DFV-A_1_B_4_ constructions. (b) Assembly schematics illustrating monomer stoichiometric ratios and incubation conditions.

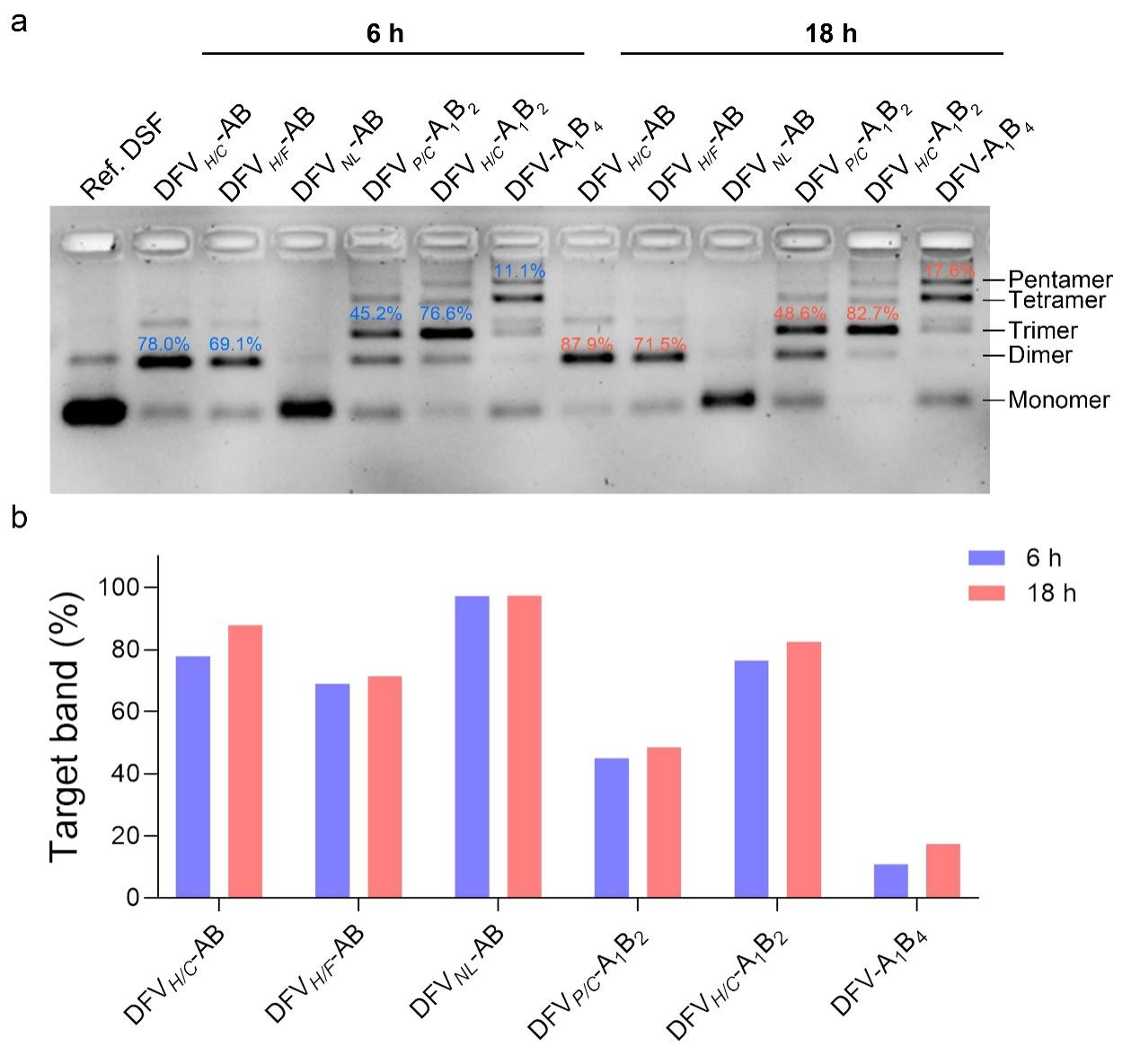

**Figure S17. Yield of DFV nanoreactors after incubation for 6 h and 18 h.**
(a) SDS-AGE results for DFV*_H/C_*-AB, DFV*_H/F_*-AB, DFV*_NL_*-AB, DFV*_P/C_*-A_1_B_2_, DFV*_H/C_*-A_1_B_2_, and DFV-A_1_B_4_ after incubation for 6 h and 18 h at 37°C. Intensity percentage of the bands at targeted assembly orders are indicated.  (b) Quantitative comparison of target band percentages between 6 h and 18 h incubation.

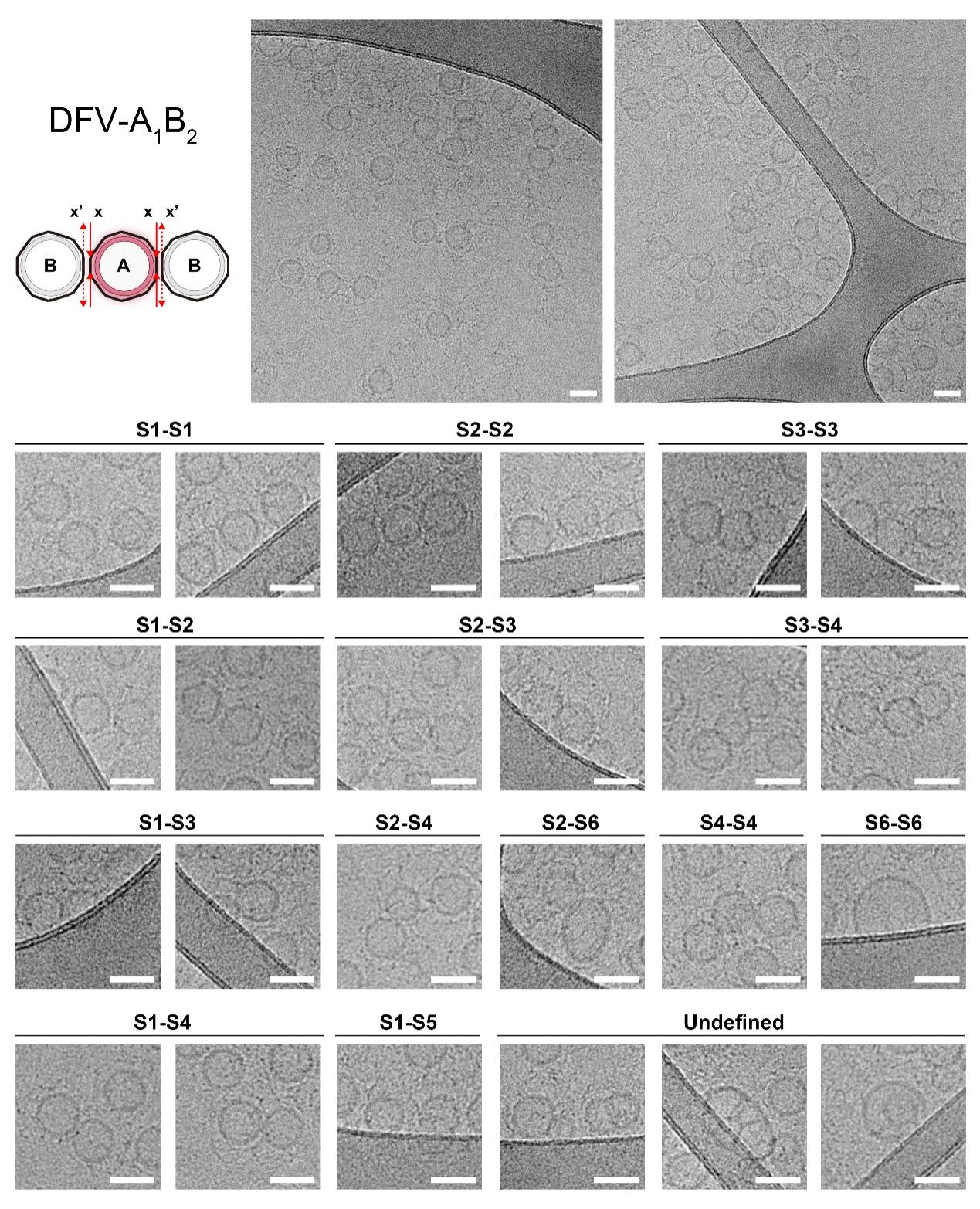

**Figure S18. Cryo-EM analysis of DFV-A_1_B_2_ trimers.** *(Top)* Representative large-scale cryo-EM images of DFV-A_1_B_2_. *(Bottom)* Gallery of classified fusion intermediates observed at the two docking windows. Each panel is labeled with the corresponding fusion stage (S1–S6, as defined in Figure 2) observed at each window. Scale bar: 50 nm.

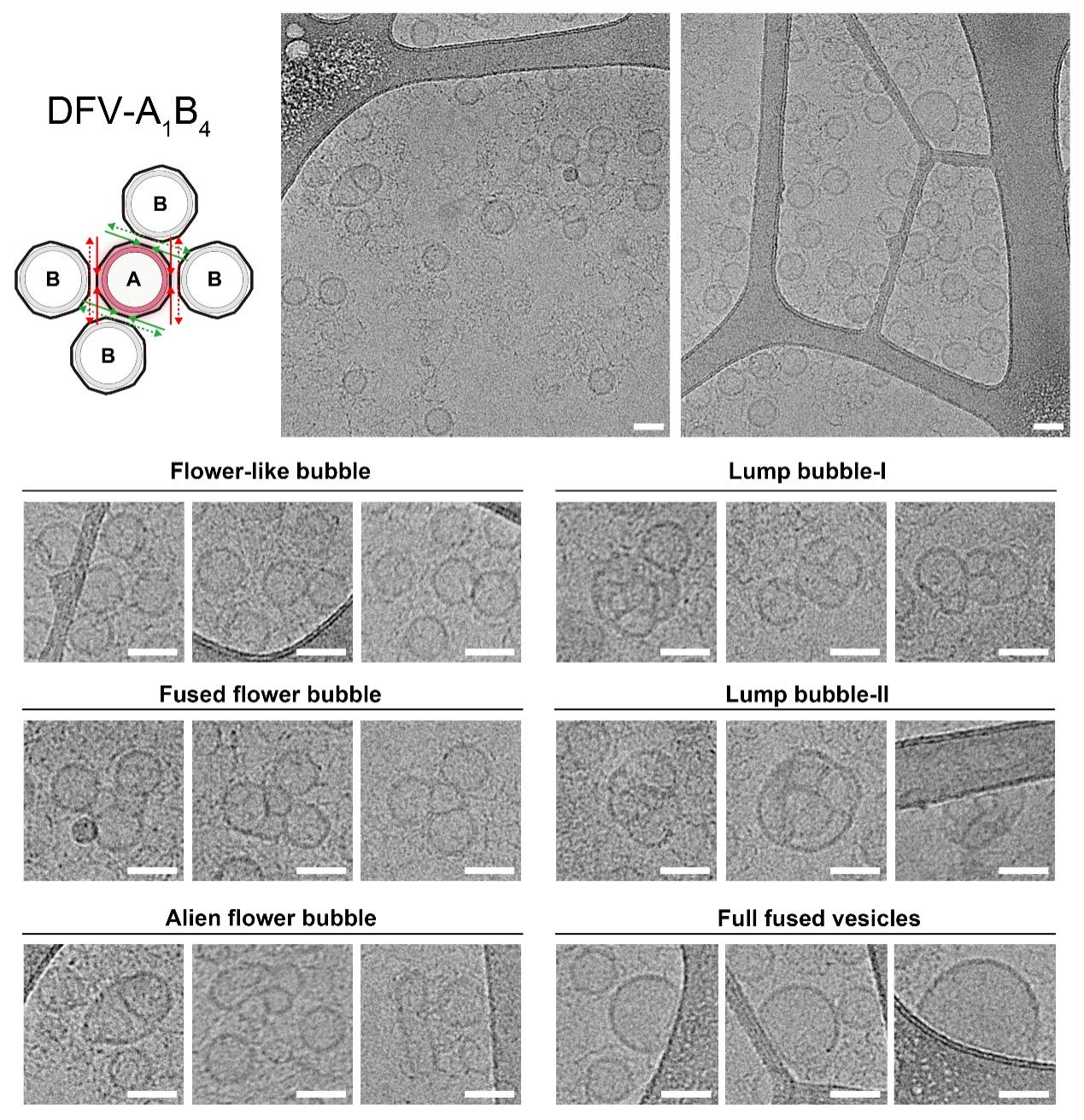

**Figure S19. Cryo-EM analysis of DFV-A_1_B_4_ pentamers.**
*(Top)* Representative large-scale Cryo-EM images of DFV-A_1_B_4_. *(Bottom)* Gallery of distinct structural morphologies observed in DFV-A_1_B_4_ assemblies, with nick names summarizing their characteristics. Scale bar: 50 nm.

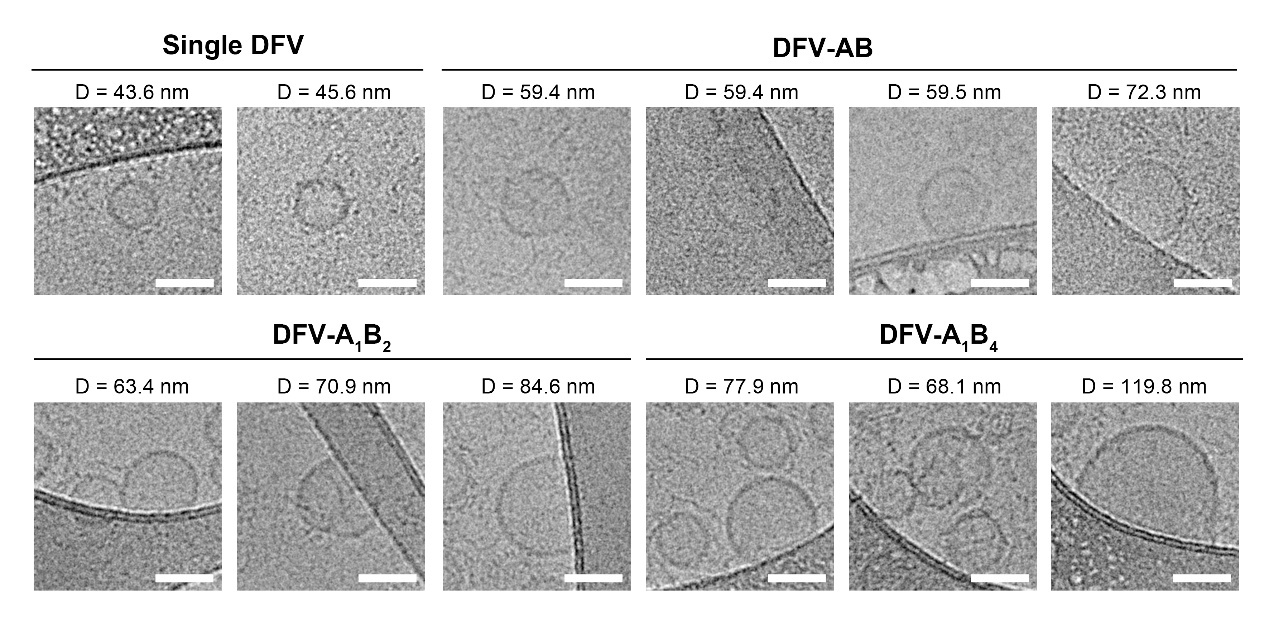

**Figure S20. Representative cryo-EM images of fully fused vesicles from DFV nanoreactors.**
Representative cryo-EM images of fully fused vesicles obtained from DFV-A_1_B_1_, DFV-A_1_B_2_, and DFV-A_1_B_4_ assemblies, along with their measured diameters in comparison with DFV monomers. Scale bar: 50 nm.

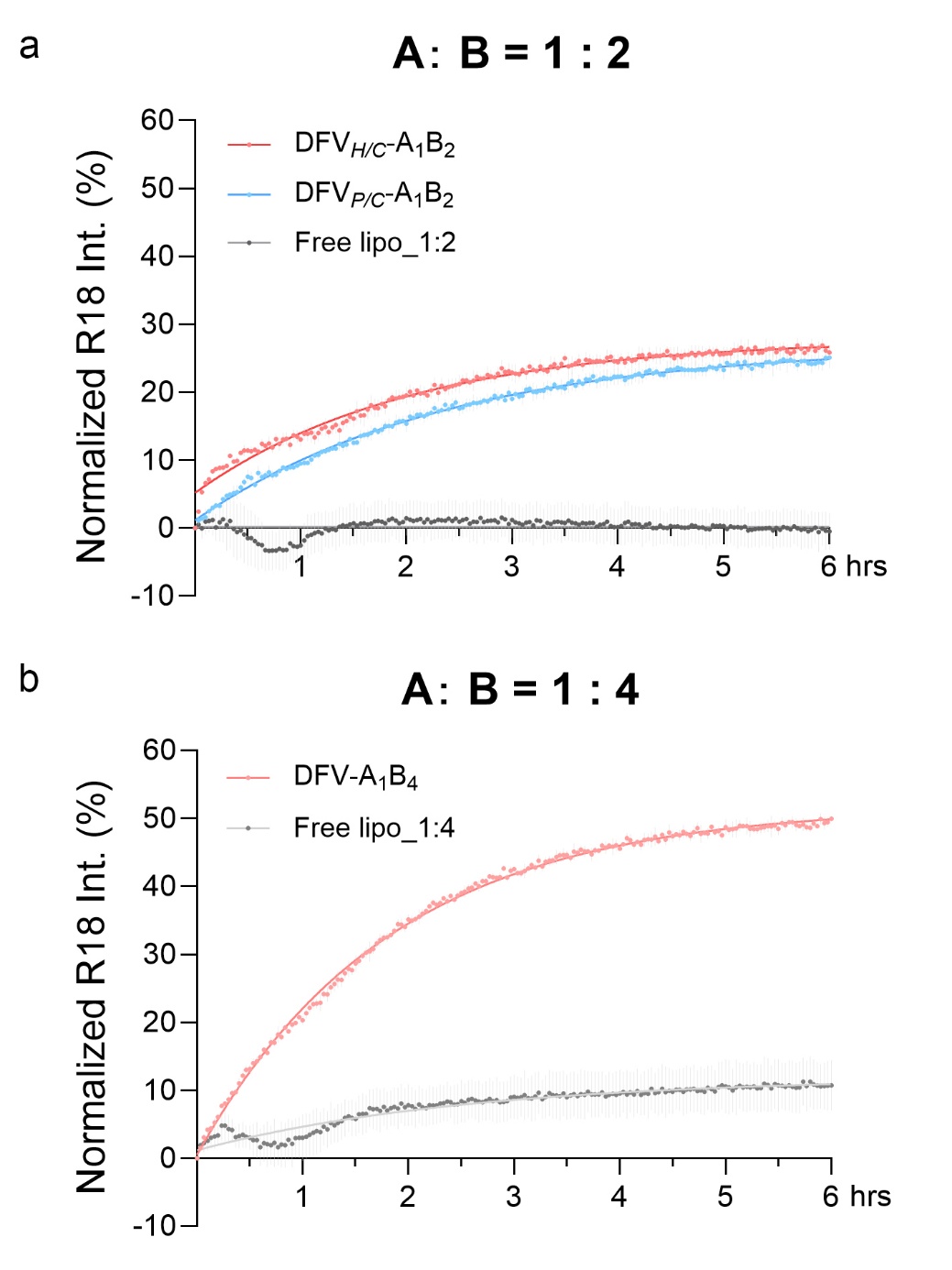

**Figure S21. Fusion efficiency of multivalent DFV assemblies.**
(a) Normalized R18 dequenching kinetics for DFV*_H/C_*-A_1_B_2_ (pink), DFV*_P/C_*-A_1_B_2_ (blue), and free liposomes (gray). (b) Fusion kinetics for DFV-A_1_B_4_ (light pink) and free liposomes (light gray) at 1:4 A:B ratio.

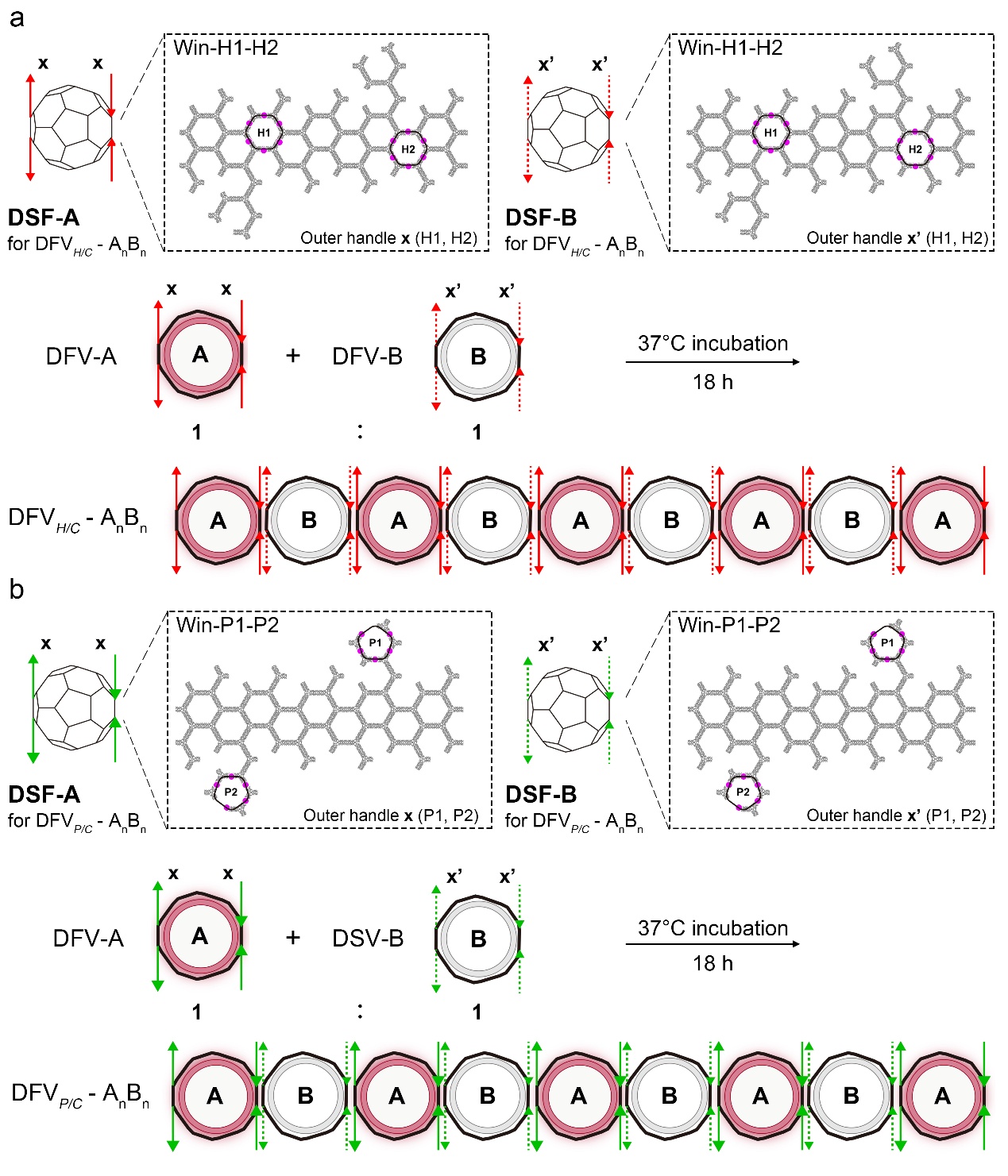

**Figure S22. Design and assembly strategy for linear DFV arrays (A_n_B_n_).**
(a) Schematic design and corresponding 2D maps showing the hexagonal docking windows used for DFV*_H/C_* - A_n_B_n_ assembly. (b) Corresponding design for DFV*_P/C_* - A_n_B_n_ using pentagonal windows. Both linear arrays were assembled by incubating DFV-A and DFV-B at a 1:1 molar ratio (37°C, 18 h).

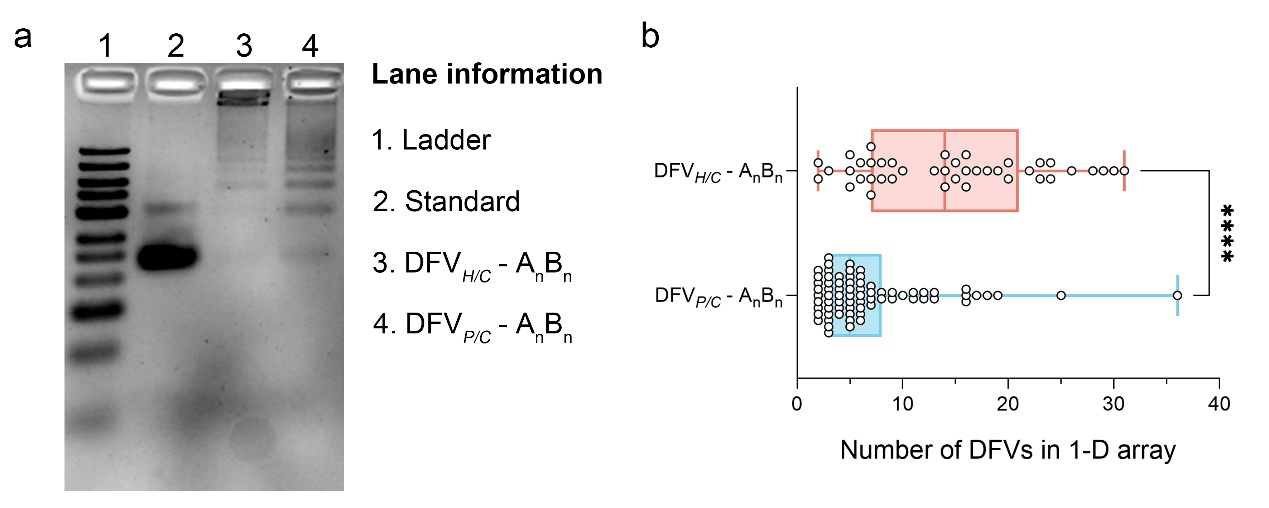

**Figure S23.** **Assembly and length distribution of linear DFV arrays (A_n_B_n_).**
(a) SDS-AGE image of the DFV*_H/C_* - A_n_B_n_ and DFV*_P/C_* - A_n_B_n_ (b) Length distribution quantifying the number of DFV units per linear array for hexagonal (*n* = 41, mean = 14.5 units) and pentagonal (*n* = 72, mean = 7.0 units) configurations.

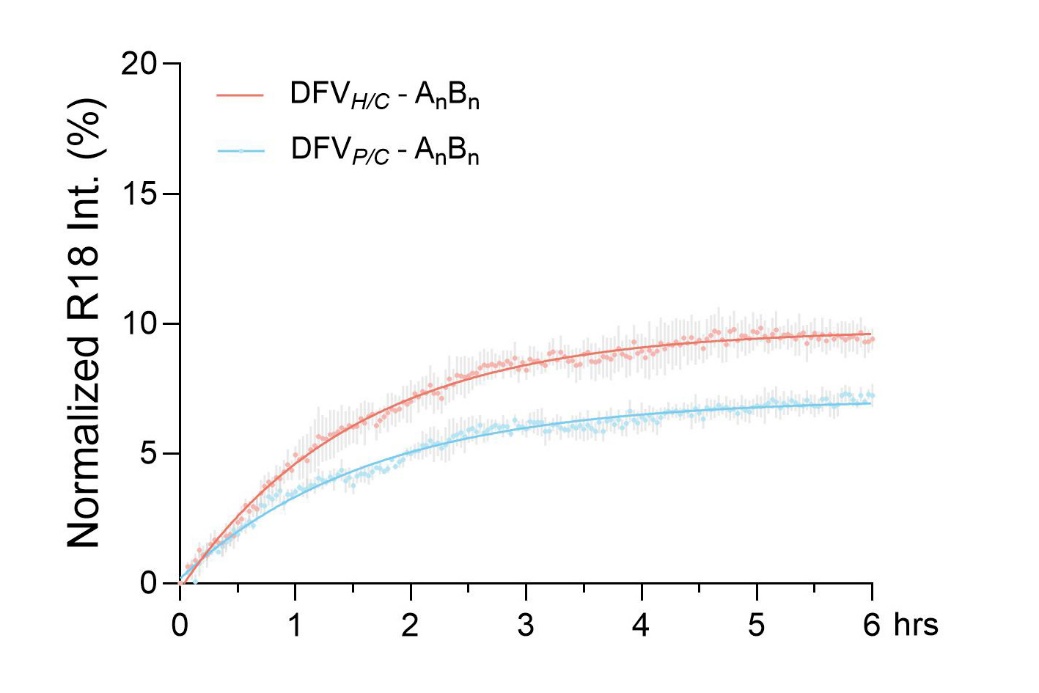

**Figure S24. Fusion kinetics of linear DFV arrays (A_n_B_n_).**
Normalized R18 dequenching traces for DFV*_H/C_* - A_n_B_n_ (light pink) and DFV*_P/C_* - A_n_B_n_ (light blue) over 6 hours.

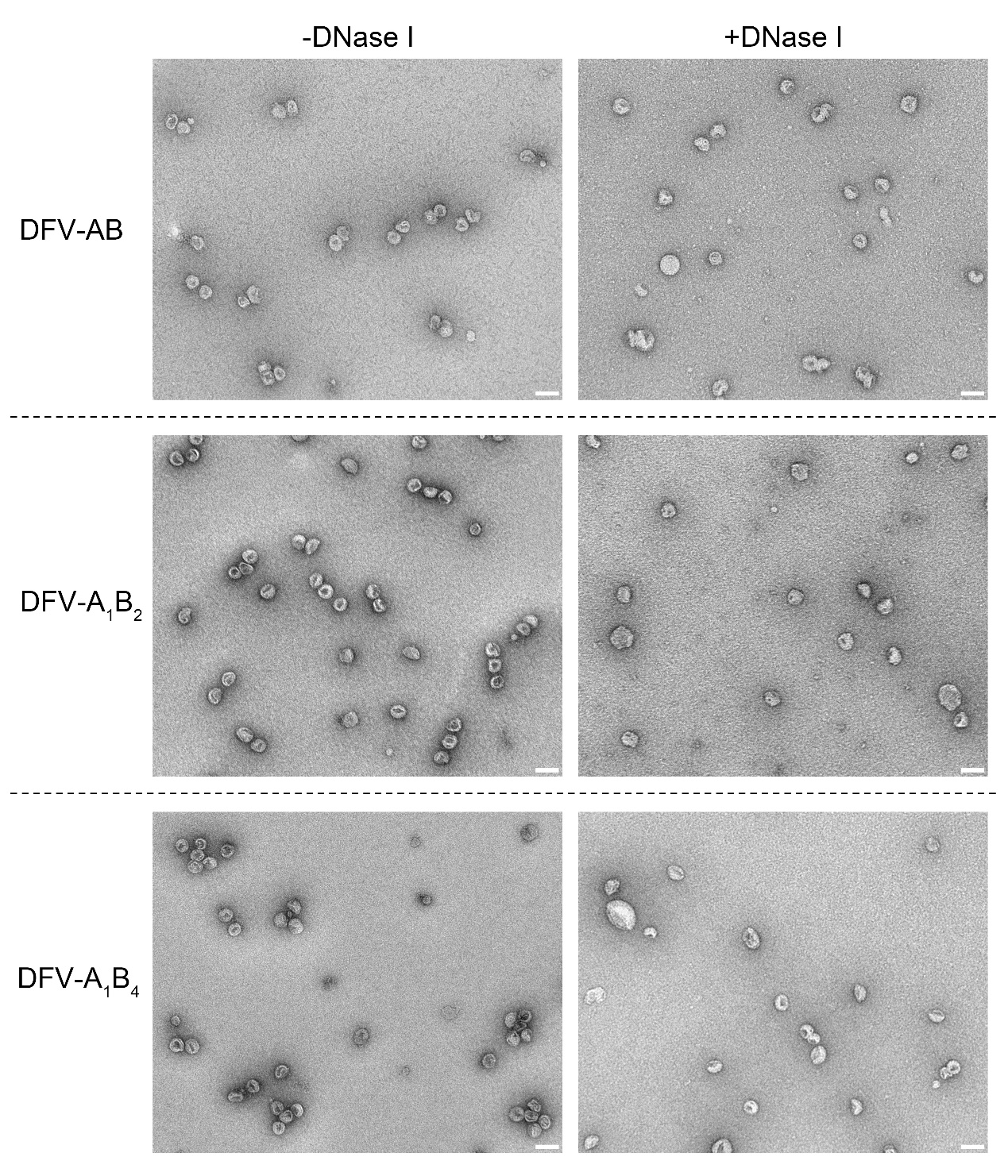

**Figure S25. nsTEM images of DFV assemblies before and after framework digestion.**
Representative nsTEM images of DFV-AB, DFV-A_1_B_2_, and DFV-A1B4 before (–DNase I) and after (+DNase I) framework digestion. Scale bar: 100 nm.

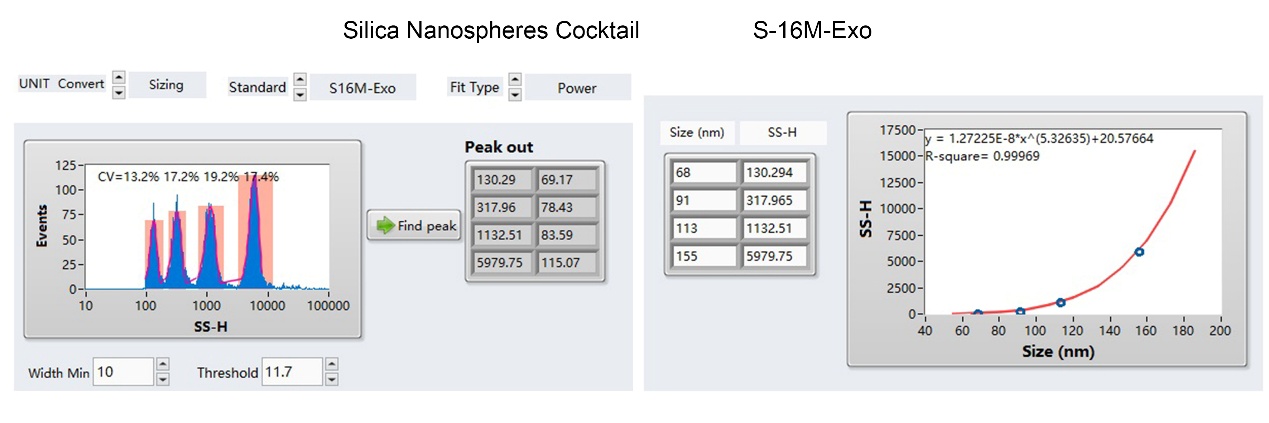

**Figure S26. nFCM size calibration using silica nanosphere standards.**
Size distribution (SSC-H signal) of a silica nanosphere cocktail (S-16M-Exo) containing four distinct sizes: 68, 91, 113, and 155 nm.

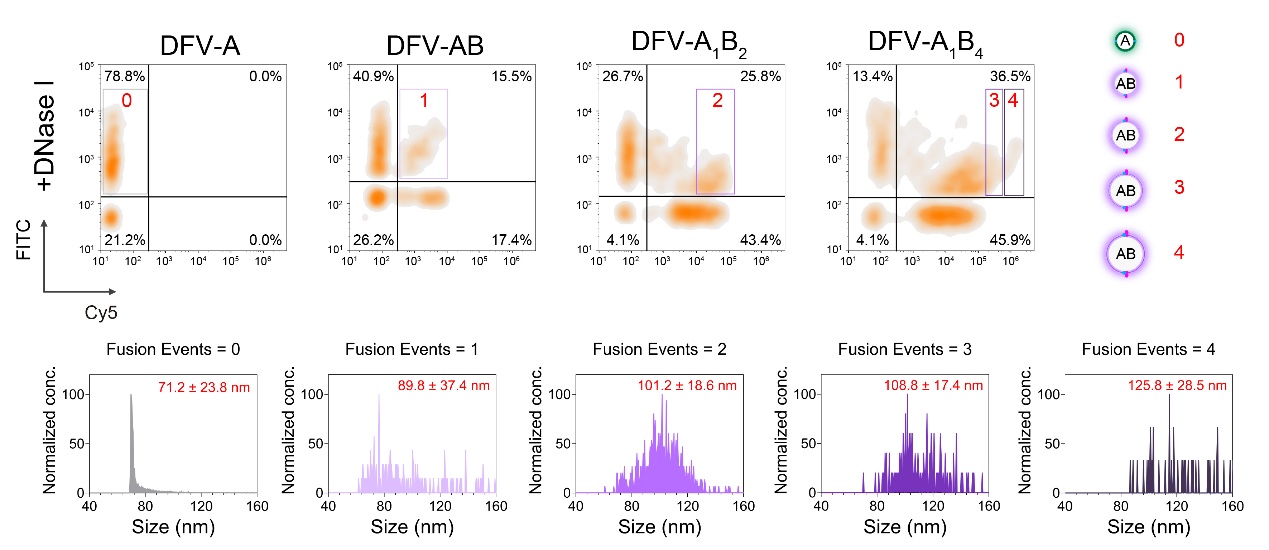

**Figure S27. Gating and size analysis of post-fusion vesicles by nFCM.** 
Following DNase I digestion of the DNA framework, nFCM analysis was performed on DFV-A, DFV-AB, DFV-A_1_B_2_, and DFV-A_1_B_4_. Subpopulations corresponding to different fusion events (unfused and fusion 1-4) were gated based on FITC/Cy5 fluorescence, and their hydrodynamic diameters (as presented in **Figure 4e**, main text) were quantified using the silica nanosphere standard curve.

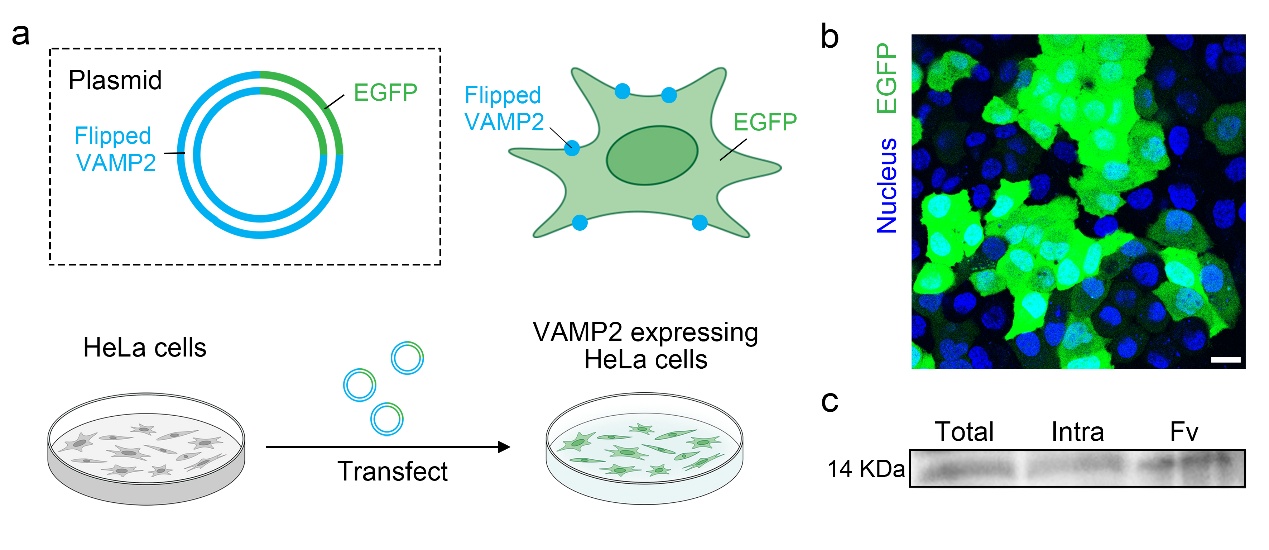

**Figure S28.  Expression and plasma membrane localization of flipped VAMP2-EGFP in HeLa cells.** (a) Schematic of the plasmid construct encoding EGFP-tagged flipped VAMP2. (b) Fluorescence micrograph of transfected HeLa cells showing EGFP expression. Scale bar, 20 µm. (c) Surface biotinylation assay analyzed by immunoblotting. Total: total cell lysate; Intra: intracellular fraction; Fv: biotinylated flipped VAMP2 isolated from the cell surface.

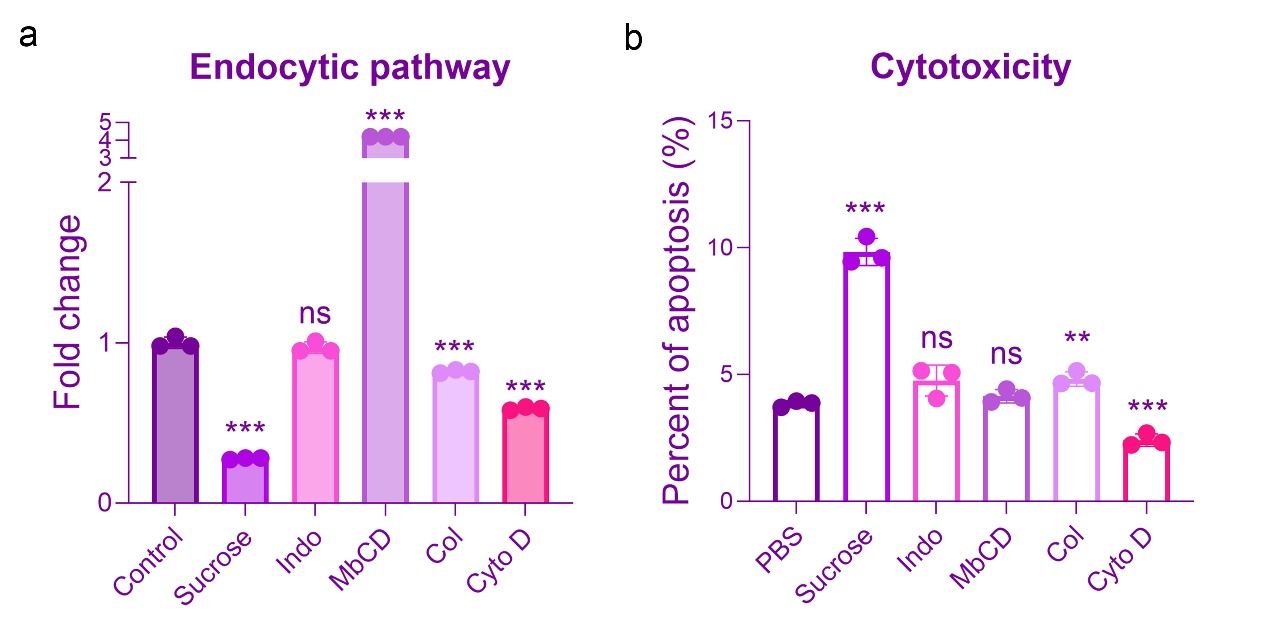

**Figure S29. Endocytosis inhibitor screening identifies cytochalasin D for cell surface-located fusion.**
(a) Relative endocytosis efficiency in HeLa cells pretreated with inhibitors targeting distinct pathways: sucrose (clathrin-mediated), indomethacin (caveolin-mediated), methyl-β-cyclodextrin (lipid raft-mediated), colchicine (macropinocytosis), and cytochalasin D (F-actin mediated). (b) Apoptosis-based cytotoxicity assessment of the inhibitor treatments. Data show mean ± SD; ns, non-significant, **P* < 0.05, ***P* < 0.01, ****P* < 0.001. Cytochalasin D was selected for subsequent experiments based on its potent inhibition and low cytotoxicity, while sucrose was used in parallel (see Figure S30 below) to support the conclusions.

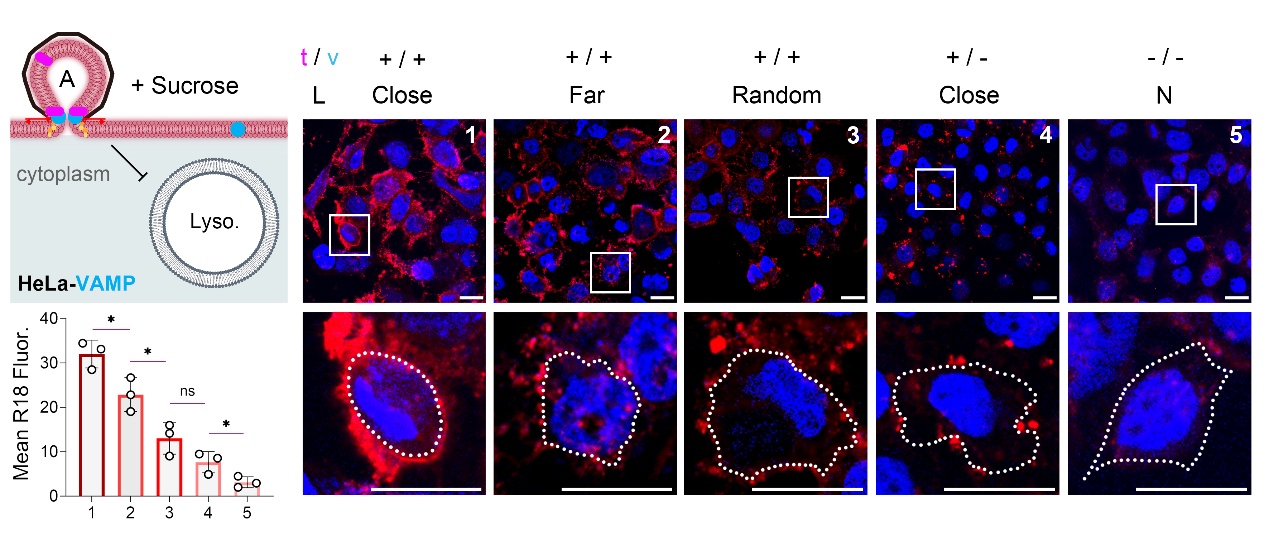

**Figure S30. Confocal microscopy images of cells incubated with DFVs under sucrose-treated conditions.** 
R18 signal (red) reports fusion extent and Hoechst (blue) stains nuclei. Quantification of R18 fluorescence intensity is shown on the left. Data are expressed as mean ± SD; ns, non-significant, **P* < 0.05.

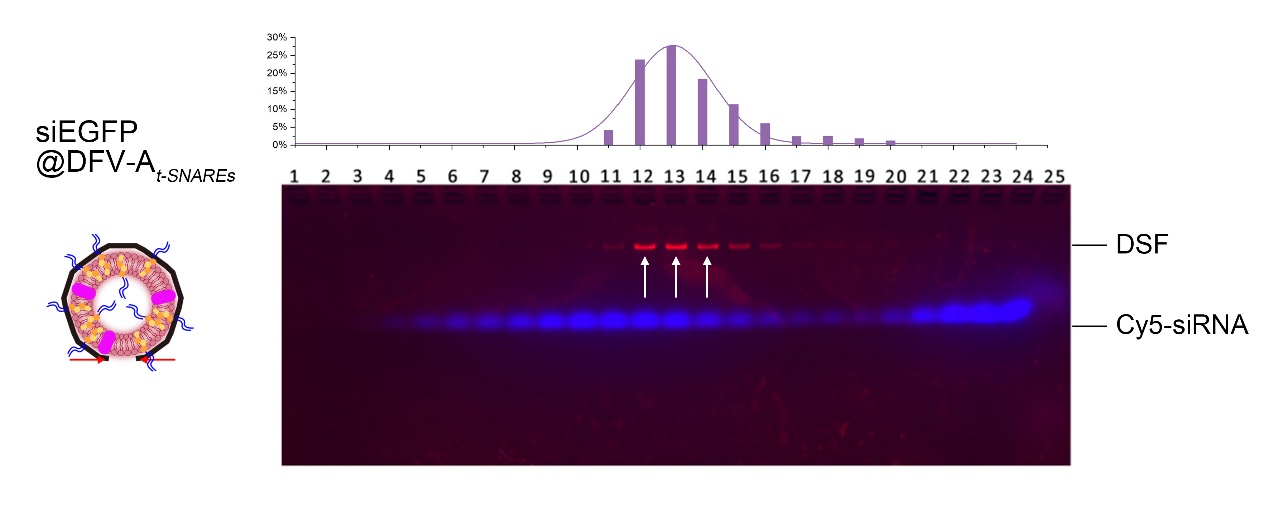

**Figure S31. Loading and purification of siEGFP in DFV-A*_t-SNARE_*.** 
SDS-AGE analysis of siEGFP-loaded DFV-A*_t-SNARE_* after density gradient purification. Peak fractions (F12-F14, white arrows) containing intact siRNA-loaded DFVs were collected for subsequent experiments.

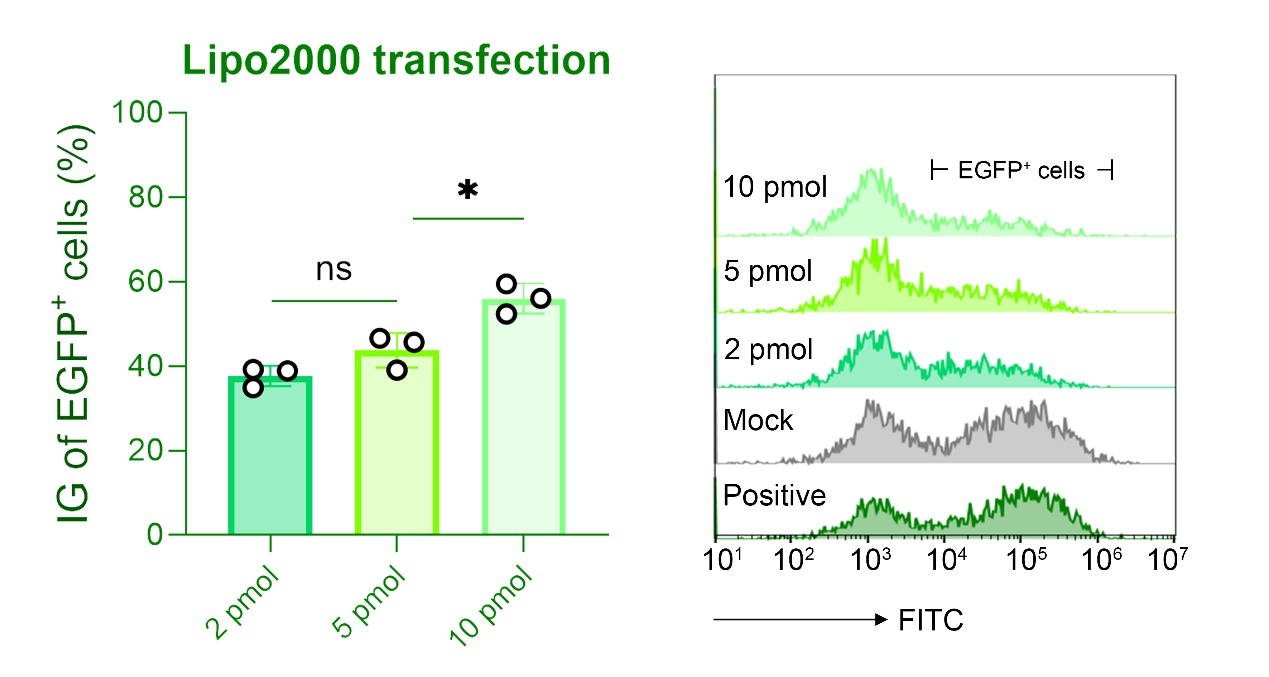

**Figure S32. Lipo2000 transfection efficiency.**
Flow cytometry analysis of EGFP inhibition grade (IG) in HeLa-EGFP-VAMP2 cells transfected with increasing doses of siEGFP (2, 5, and 10 pmol) using Lipo2000. Mean ± SD; ns, not significant, **P* < 0.05.

3. **Supplementary Tables**

**Table S1.** DNA strand sequences

| **Inner handle for nucleation seed decoration** | |
| --- | --- |
| Name | Sequence (5'-3') |
| orange-In-006 | AGGTAAAGATAACAGCT**T**TAACATTCCTAACTTCTCATA |
| orange-In-007 | GCTGATAAATTTTCACA**T**TAACATTCCTAACTTCTCATA |
| orange-In-008 | GTCTGGAGCAAGGCTGC**T**TAACATTCCTAACTTCTCATA |
| orange-In-009 | ATTCCATATAACAAGAAAAGCCCCA**T**TAACATTCCTAACTTCTCATA |
| orange-In-010 | CCTTTATTTCCTGGATA**T**TAACATTCCTAACTTCTCATA |
| orange-In-021 | CCGGCGAACGTGGAATCCTGTTTGA**T**TAACATTCCTAACTTCTCATA |
| orange-In-023 | CACACGACCAGTACAAGGCGATTAA**T**TAACATTCCTAACTTCTCATA |
| orange-In-024 | ATCTAAAGCATCATTTGTTAAATCA**T**TAACATTCCTAACTTCTCATA |
| orange-In-025 | ATTATCATTTTGCGAGAATGACCAT**T**TAACATTCCTAACTTCTCATA |
| orange-In-041 | AATCAAGTTTTAGGAAT**T**TAACATTCCTAACTTCTCATA |
| orange-In-042 | AATAATCGGCCACACCC**T**TAACATTCCTAACTTCTCATA |
| orange-In-045 | GCAACAGGAATTAGTTA**T**TAACATTCCTAACTTCTCATA |
| orange-In-046 | CTCCGGCTTATTGAATG**T**TAACATTCCTAACTTCTCATA |
| orange-In-047 | CCAGCAGAAGATATATG**T**TAACATTCCTAACTTCTCATA |
| orange-In-050 | AATATAATCCTGAATTACAGGTAGA**T**TAACATTCCTAACTTCTCATA |
| orange-In-066 | CTTACCGAAGCCCCCTGTTTATCAA**T**TAACATTCCTAACTTCTCATA |
| orange-In-067 | TAGGATTAGCGGGAAACACCGGAAT**T**TAACATTCCTAACTTCTCATA |
| orange-In-068 | AAAGGAATTGCGAGATAGCTTAGAT**T**TAACATTCCTAACTTCTCATA |
| orange-In-070 | CGTTTTAGCGTAAGAAC**T**TAACATTCCTAACTTCTCATA |
| orange-In-081 | GCCAGCAAAATTACCAA**T**TAACATTCCTAACTTCTCATA |
| orange-In-082 | AATACCCAAAAGAACCAGAGCCGCC**T**TAACATTCCTAACTTCTCATA |
| orange-In-083 | ACCGCCTCCCTCAGGAG**T**TAACATTCCTAACTTCTCATA |
| orange-In-084 | CGGTCATAGCCTTTCGA**T**TAACATTCCTAACTTCTCATA |
| orange-In-085 | GCACCGTAATATCCGCG**T**TAACATTCCTAACTTCTCATA |
| **Nucleation seed sequence** | |
| In'-Chol | TATGAGAAGTTAGGAATGTTA/3Chol/ |
| **DNA linker strands for DFV*_H/C_*-AB** | |
| x-orange-040 | CTTCTTCACGTTTACACGGCCTTCAATACCTTTAAGCAAACTCCAATAGATTAGAGCC |
| x-orange-048 | CTTCTTCACGTTTACACGGCCTTCAATACCTTTGAAGATGATGGGTTATC |
| x-orange-049 | CTTCTTCACGTTTACACGGCCTTCAATACCTTTTTAGAAGTATAGAAATT |
| x-orange-063 | CTTCTTCACGTTTACACGGCCTTCAATACCTTTAGACTTTTTCATGCAGAGGCGAATT |
| x-orange-064 | CTTCTTCACGTTTACACGGCCTTCAATACCTTTGGCTGGCTGACCTATATACAGTAAC |
| x-orange-069 | CTTCTTCACGTTTACACGGCCTTCAATACCTTTTAAAACACTCATCGGATTCGCCTGA |
| yellow-x'-022 | AAGACGGAGTGAGAAGTTTTGGTATTGAAGGCCGTGTAAACGTGAAGAAG |
| yellow-x'-028 | CATACGAGCCCGATTAATTTGGTATTGAAGGCCGTGTAAACGTGAAGAAG |
| yellow-x'-029 | GCTTCTAATGTCTGTCCTTTGGTATTGAAGGCCGTGTAAACGTGAAGAAG |
| yellow-x'-043 | TACCGACAAGCCAGTAATTTGGTATTGAAGGCCGTGTAAACGTGAAGAAG |
| yellow-x'-044 | CTGAGTAGATGATTAGTTTTGGTATTGAAGGCCGTGTAAACGTGAAGAAG |
| yellow-x'-052 | TAATAAGTTTTTAACAATTTGGTATTGAAGGCCGTGTAAACGTGAAGAAG |
| **DSF tracking strand** | |
| blue-031-TAMRA | GAACAGCCCCCGATTTTTTTAGAGC/3TAMRA/ |
| blue-037-TAMRA | CTCACAGGAACGGTATTTTCGCCAG/3TAMRA/ |
| blue-043-TAMRA | GGTGGGGATTATTTATTTTCATTGG/3TAMRA/ |
| blue-049-TAMRA | AAATGCCACGCTGAGTTTTAGCCAG/3TAMRA/ |
| blue-055-TAMRA | CCGAAACGTTATTAATTTTTTTTAA/3TAMRA/ |
| **DNA linker strands for DFV*_P/C_*-AB** | |
| x-orange-001 | CTTCTTCACGTTTACACGGCCTTCAATACCTTTACATCCAATAAATAGTCAAATCACC |
| x-orange-002 | CTTCTTCACGTTTACACGGCCTTCAATACCTTTGCTGAAAAGGTGGGAGAGATCTACA |
| x-orange-003 | CTTCTTCACGTTTACACGGCCTTCAATACCTTTTGGTCAATAACCTGTAAAACTAGCA |
| x-orange-004 | CTTCTTCACGTTTACACGGCCTTCAATACCTTTGCGAACGAGTATTATGA |
| x-orange-005 | CTTCTTCACGTTTACACGGCCTTCAATACCTTTTTAAGCAATAAAGCCTCATATATTT |
| yellow-x'-086 | GTCACCAATCGATTGAGTTTGGTATTGAAGGCCGTGTAAACGTGAAGAAG |
| yellow-x'-087 | CGACTTGAGTAAAGGTGTTTGGTATTGAAGGCCGTGTAAACGTGAAGAAG |
| yellow-x'-088 | ACCACCCTCTACATACATTTGGTATTGAAGGCCGTGTAAACGTGAAGAAG |
| yellow-x'-089 | CACCGGAACACCACGGATTTGGTATTGAAGGCCGTGTAAACGTGAAGAAG |
| yellow-x'-090 | TGTAGCGCGATATGGTTTTTGGTATTGAAGGCCGTGTAAACGTGAAGAAG |
| **DNA linker strands for DFV*_H/F_*-AB** | |
| yellow-f-040 | AAGCAAACTCCAATAGATTAGAGCCTAAATTATCTACCACAACTCAC |
| yellow-f-048 | GAAGATGATGGGTTATCTAAATTATCTACCACAACTCAC |
| yellow-f-049 | TTAGAAGTATAGAAATTTAAATTATCTACCACAACTCAC |
| yellow-f-063 | AGACTTTTTCATGCAGAGGCGAATTTAAATTATCTACCACAACTCAC |
| yellow-f-064 | GGCTGGCTGACCTATATACAGTAACTAAATTATCTACCACAACTCAC |
| yellow-f-069 | TAAAACACTCATCGGATTCGCCTGATAAATTATCTACCACAACTCAC |
| yellow-f'-022 | AAGACGGAGTGAGAAGT**T**GTGAGTTGTGGTAGATAATTT |
| yellow-f'-028 | CATACGAGCCCGATTAA**T**GTGAGTTGTGGTAGATAATTT |
| yellow-f'-029 | GCTTCTAATGTCTGTCC**T**GTGAGTTGTGGTAGATAATTT |
| yellow-f'-043 | TACCGACAAGCCAGTAA**T**GTGAGTTGTGGTAGATAATTT |
| yellow-f'-044 | CTGAGTAGATGATTAGT**T**GTGAGTTGTGGTAGATAATTT |
| yellow-f'-052 | TAATAAGTTTTTAACAA**T**GTGAGTTGTGGTAGATAATTT |
| **DNA linker strands for DFV-A_1_B_2_ and DFV-A_1_B_4_** | |
| x-orange-040、048、049、063、064、069 | Listed in DNA linker strands for DFV-AB-hex |
| x-orange-022 | CTTCTTCACGTTTACACGGCCTTCAATACC**TTT**GTTTTTATAATCAGTTACCTCGATA |
| x-orange-028 | CTTCTTCACGTTTACACGGCCTTCAATACC**TTT**AGGGATTTTAGACAATTCCACACAA |
| x-orange-029 | CTTCTTCACGTTTACACGGCCTTCAATACC**TTT**ATCACGCAAATTACAAAATAACCCC |
| x-orange-043 | CTTCTTCACGTTTACACGGCCTTCAATACC**TTT**TCCTCGTTAGTATAAAG |
| x-orange-044 | CTTCTTCACGTTTACACGGCCTTCAATACC**TTT**ACGCTCAACACACTTGC |
| x-orange-052 | CTTCTTCACGTTTACACGGCCTTCAATACC**TTT**CGCCAACATGTAAGGAGTGTACTGG |
| yellow-x'-022、028、029、043、044、052 | Listed in DNA linker strands for DFV-AB-hex |
| y-orange-001 | CCGTTCACAACGTCCATCTTGATTTCCATC**T**ACATCCAATAAATAGTCAAATCACC |
| y-orange-002 | CCGTTCACAACGTCCATCTTGATTTCCATC**T**GCTGAAAAGGTGGGAGAGATCTACA |
| y-orange-003 | CCGTTCACAACGTCCATCTTGATTTCCATC**T**TGGTCAATAACCTGTAAAACTAGCA |
| y-orange-004 | CCGTTCACAACGTCCATCTTGATTTCCATC**T**GCGAACGAGTATTATGA |
| y-orange-005 | CCGTTCACAACGTCCATCTTGATTTCCATC**T**TTAAGCAATAAAGCCTCATATATTT |
| y-orange-086 | CCGTTCACAACGTCCATCTTGATTTCCATC**T**GGAGGGAAGGTAAAAGGCCGGAAAC |
| y-orange-087 | CCGTTCACAACGTCCATCTTGATTTCCATC**T**CTCCTTATTACCGTCAC |
| y-orange-088 | CCGTTCACAACGTCCATCTTGATTTCCATC**T**TAAAGGTGGCAACACCCTCAGAGCC |
| y-orange-089 | CCGTTCACAACGTCCATCTTGATTTCCATC**T**ATAAGTTTATTTTCATAATCAAAAT |
| y-orange-090 | CCGTTCACAACGTCCATCTTGATTTCCATC**T**TACCAGCGCCAAATTAGCGTCAGAC |
| yellow-y'-001 | ATCAATATGTAGCATTA**T**GATGGAAATCAAGATGGACGTTGTGAACGG |
| yellow-y'-002 | AAGGCTATCGGGCGCGA**T**GATGGAAATCAAGATGGACGTTGTGAACGG |
| yellow-y'-003 | TGTCAATCATTCGCAAA**T**GATGGAAATCAAGATGGACGTTGTGAACGG |
| yellow-y'-004 | CCCTGTAATCGGTTGTA**T**GATGGAAATCAAGATGGACGTTGTGAACGG |
| yellow-y'-005 | TAAATGCAATAGCAAAA**T**GATGGAAATCAAGATGGACGTTGTGAACGG |
| **DNA linker strands for DFV*_H/C_* - A_n_B_n_** | |
| x-orange-040、048、049、063、064、069 | Listed in DNA linker strands for DFV*_H/C_*-AB |
| yellow-x-022 | AAGACGGAGTGAGAAGT**TTT**CTTCTTCACGTTTACACGGCCTTCAATACC |
| yellow-x-028 | CATACGAGCCCGATTAA**TTT**CTTCTTCACGTTTACACGGCCTTCAATACC |
| yellow-x-029 | GCTTCTAATGTCTGTCC**TTT**CTTCTTCACGTTTACACGGCCTTCAATACC |
| yellow-x-043 | TACCGACAAGCCAGTAA**TTT**CTTCTTCACGTTTACACGGCCTTCAATACC |
| yellow-x-044 | CTGAGTAGATGATTAGT**TTT**CTTCTTCACGTTTACACGGCCTTCAATACC |
| yellow-x-052 | TAATAAGTTTTTAACAA**TTT**CTTCTTCACGTTTACACGGCCTTCAATACC |
| x'-orange-040 | GGTATTGAAGGCCGTGTAAACGTGAAGAAG**TTT**AAGCAAACTCCAATAGATTAGAGCC |
| x'-orange-048 | GGTATTGAAGGCCGTGTAAACGTGAAGAAG**TTT**GAAGATGATGGGTTATC |
| x'-orange-049 | GGTATTGAAGGCCGTGTAAACGTGAAGAAG**TTT**TTAGAAGTATAGAAATT |
| x'-orange-063 | GGTATTGAAGGCCGTGTAAACGTGAAGAAG**TTT**AGACTTTTTCATGCAGAGGCGAATT |
| x'-orange-064 | GGTATTGAAGGCCGTGTAAACGTGAAGAAG**TTT**GGCTGGCTGACCTATATACAGTAAC |
| x'-orange-069 | GGTATTGAAGGCCGTGTAAACGTGAAGAAG**TTT**TAAAACACTCATCGGATTCGCCTGA |
| yellow-x'-022、028、029、043、044、052 | Listed in DNA linker strands for DFV*_H/C_*-AB |
| **DNA linker strands for DFV*_P/C_* - A_n_B_n_** | |
| x-orange-001、002、003、004、005 | Listed in DNA linker strands for DFV*_P/C_*-AB |
| yellow-x-086 | GTCACCAATCGATTGAG**TTT**CTTCTTCACGTTTACACGGCCTTCAATACC |
| yellow-x-087 | CGACTTGAGTAAAGGTG**TTT**CTTCTTCACGTTTACACGGCCTTCAATACC |
| yellow-x-088 | ACCACCCTCTACATACA**TTT**CTTCTTCACGTTTACACGGCCTTCAATACC |
| yellow-x-089 | CACCGGAACACCACGGA**TTT**CTTCTTCACGTTTACACGGCCTTCAATACC |
| yellow-x-090 | TGTAGCGCGATATGGTT**TTT**CTTCTTCACGTTTACACGGCCTTCAATACC |
| x'-orange-001 | GGTATTGAAGGCCGTGTAAACGTGAAGAAG**TTT**ACATCCAATAAATAGTCAAATCACC |
| x'-orange-002 | GGTATTGAAGGCCGTGTAAACGTGAAGAAG**TTT**GCTGAAAAGGTGGGAGAGATCTACA |
| x'-orange-003 | GGTATTGAAGGCCGTGTAAACGTGAAGAAG**TTT**TGGTCAATAACCTGTAAAACTAGCA |
| x'-orange-004 | GGTATTGAAGGCCGTGTAAACGTGAAGAAG**TTT**GCGAACGAGTATTATGA |
| x'-orange-005 | GGTATTGAAGGCCGTGTAAACGTGAAGAAG**TTT**TTAAGCAATAAAGCCTCATATATTT |
| yellow-x'-086、087、088、089、090 | Listed in DNA linker strands for DFV*_P/C_*-AB |

**Table S2.** Lipid information.

| **Abbreviation** | **Full name of lipids** | **M.W.** |
| --- | --- | --- |
| DOPC | 1,2-dioleoyl-sn-glycero-3-phosphocholine | 786.113 |
| DOPS | 1,2-Dioleoyl-sn-glycero-3-phospho-l-serine | 810.025 |
| DOPE | 1,2-dioleoyl-sn-glycero-3-phosphoethanolamine | 744.034 |
| PI(4,5)P2 | phosphatidylinositol 4,5-bisphosphate | 1074.158 |
| PEG-2k-DOPE | 1,2-dioleoyl-sn-glycero-3-phosphoethanolamine-N-[methoxy(polyethylene glycol)-2000] | 2801.465 |
| R18 | octadecyl Rhodamine B chloride | 731.4999 |
| FITC-Chol | FITC modified cholesterol | 776.05 |
| Cy5-Chol | Cy5 modified cholesterol | 1043.46 |
| NBD-DOPE | 1,2-dioleoyl-sn-glycero-3-phosphoethanolamine-N-(7-nitro-2-1,3-benzoxadiazol-4-yl) | 872.09 |
| Egg Liss Rhod-PE | L-α-Phosphatidylethanolamine-N-(lissamine rhodamine B sulfonyl) | 1275.678 |

**Table S3.** Lipid compositions for construction of liposomes and Buffer ingredients. Numeric values refer to molar percentages and ratios.

|  | **DOPC** | **DOPS** | **DOPE** | **Fluorescent lipid** | **PIP2** | **PEG-2k-DOPE** |
| --- | --- | --- | --- | --- | --- | --- |
| Clear Lipid | 76% | 10% | 10% | 0% | 2% | 2% |
| R18 Lipid | 73% | 10% | 10% | 5% (R18) | 0% | 2% |
| Cy5 Lipid | 74% | 10% | 10% | 2% (Cy5) | 2% | 2% |
| FITC Lipid | 76% | 10% | 10% | 2% (FITC) | 0% | 2% |
| NBD&Rho Lipid | 75% | 10% | 10% | 1.5% (NBD&Rho) | 0% | 2% |

|  | **Lipid compositions** | **OG (w/v)** | **Protein:Lipid** |
| --- | --- | --- | --- |
| t-SNARE Clear liposome | Clear Lipid | 1% | 400:1 |
| t-SNARE Cy5 liposome | Cy5 Lipid | 1% | 400:1 |
| VAMP2 R18 liposome | R18 Lipid | 1% | 200:1 |
| VAMP2 FITC liposome | FITC Lipid | 1% | 200:1 |
| VAMP2 NBD&Rho liposome | NBD&Rho Lipid | 1% | 200:1 |

|  | **HEPES** | **KCl** | **MgCl_2_** | **pH** |
| --- | --- | --- | --- | --- |
| 1× Hydration Buffer (HB) | 25 mM | 400 mM | 10 mM | 7.0 |

**Table S4.** siEGFP three-strand bridge sequences.

| Name | Sequence (5'-3') |
| --- | --- |
| Chol-anchor | /5Chol/TTTACCTACTAACATAATCATCAC/ |
| siEGFP sense | GCCACAACGUCUAUAUCAUdTdT |
| anti-anchor-antisense | AUGAUAUAGACGUUGUGGCGUGAUGAUUAUGUUAGUAGGU-dTdT |
